# *Dolosigranulum pigrum* cooperation and competition in human nasal microbiota

**DOI:** 10.1101/678698

**Authors:** Silvio D. Brugger, Sara M. Eslami, Melinda M. Pettigrew, Isabel F. Escapa, Matthew T. Henke, Yong Kong, Katherine P. Lemon

## Abstract

**Background:** Multiple epidemiological studies identify *Dolosigranulum pigrum* as a candidate beneficial bacterium based on its positive association with health, including negative associations with nasal/nasopharyngeal colonization by the pathogenic species *Staphylococcus aureus* and *Streptococcus pneumoniae*.

**Results:** Using a multipronged approach to gain new insights into *D. pigrum* function, we observed phenotypic interactions and predictions of genomic capacity that support a role for microbe-microbe interactions involving *D. pigrum* in shaping the composition of human nasal microbiota. We identified *in vivo* community-level and *in vitro* phenotypic cooperation by specific nasal *Corynebacterium* species. Also, *D. pigrum* inhibited *S. aureus* growth *in vitro*. Whereas, robust inhibition of *S. pneumoniae* required both *D. pigrum* and a nasal *Corynebacterium* together, and not either alone. *D. pigrum* L-lactic-acid production was insufficient to account for these inhibitions. Genomic analysis of 11 strains revealed that *D. pigrum* has a small genome (average 1.86 Mb) and multiple predicted auxotrophies consistent with *D. pigrum* relying on its human host and cocolonizing bacteria for key nutrients. Further, the accessory genome of *D. pigrum* encoded a diverse repertoire of biosynthetic gene clusters, some of which may have a role in microbe-microbe interactions.

**Conclusions:** These new insights into *D. pigrum*’s functions advance the field from compositional analysis to genomic and phenotypic experimentation on a potentially beneficial bacterial resident of the human upper respiratory tract and lay the foundation for future animal and clinical experiments.

## Background

Colonization of the human nasal passages by *Staphylococcus aureus* or *Streptococcus pneumoniae* is a major risk factor for infection by the colonizing bacterium at a distant body site [1–5]. Interventions that reduce the prevalence of colonization also reduce the risk of infection and transmission, e.g., as in [6, 7]. *S. aureus* and *S. pneumoniae* are major human pathogens that cause significant morbidity and mortality worldwide [8–11]. There are also concerns regarding rising rates of antimicrobial resistance [12] and the potential for long-term effects of antibiotics early in life [13]. Thus, efforts have recently focused on the identification of candidate bacteria that confer colonization resistance against *S. aureus* [14–21] and *S. pneumoniae* [22–25], with particular urgency for *S. aureus* in the absence of an effective vaccine.

*Dolosigranulum pigrum* has emerged in multiple studies of the human upper respiratory tract microbiota, colonizing with or without *Corynebacterium* species, as potentially beneficial and/or protective against colonization by *S. aureus* and *S. pneumoniae* [26–50] (reviewed in [14, 51–54]). Little is known about this Gram-positive, catalase-negative, Firmicute bacterium, first described in 1993 [55]. Microbiota studies sampling either nostrils or nasopharynx show very similar results; therefore, for simplicity, we use nasal or nasal passages to denote the area inclusive of the nostrils through the nasopharynx. *D. pigrum* and *S. aureus* are inversely correlated in adult nasal microbiota [31, 41, 56]. Whereas, in pediatric nasal microbiota, *D. pigrum* and members of the genus *Corynebacterium* are overrepresented when *S. pneumoniae* is absent [26, 33, 40]. Moreover, children with *D. pigrum* colonization of the nasal passages are less likely to have acute otitis media [27, 40] and it has been speculated that *D. pigrum*-dominated microbiota profiles might be more resistant to invasive pneumococcal disease [45]. Furthermore, *D. pigrum* abundance in the nasal passages is inversely associated with wheezing and respiratory tract infections in infants [28] and abundance of *D. pigrum* with *Corynebacterium* in adults provides greater community stability in the face of pneumococcal exposure [50]. The intriguing inference from these studies that *D. pigrum* plays a beneficial role in human nasal microbiota deserves further investigation.

In contrast to the above, there are very few reports of *D. pigrum* in association with human disease [57–61]. Its frequent identification in human nasal microbiota [26, 30–32, 34–37, 39–45, 47, 48, 62–73] coupled with its rare association with infection are consistent with *D. pigrum* functioning as a commensal, and possibly as a mutualist, of humans––characteristics that support its potential for future use as a therapeutic. However, its metabolism and its interplay with other nasal bacteria remain uncharted territory. Using a multipronged approach, we have made significant advances in these areas. First, we identified specific species of candidate bacterial interactors with *D. pigrum* by analyzing nasal microbiota datasets from adults and children. Second, we used *in vitro* phenotypic assays to show that *D. pigrum* exhibits distinct interaction phenotypes with nasal *Corynebacterium* species, *S. aureus* and *S. pneumoniae*. Third, based on the genomes of 11 distinct *D. pigrum* strains, we identify key predicted functions and auxotrophies in its core genome plus a diversity of predicted biosynthetic gene clusters in its accessory genome. This critical shift to phenotypic and genomic experimentation marks a significant advance in understanding *D. pigrum*, a potential beneficially member of human nasal microbiota.

## Results

### Individual bacterial species are associated with *D. pigrum* in the nasal microbiota of both adults and children

*D. pigrum* is the only member of its genus and multiple genus-level 16S rRNA gene-based nasal microbiota studies have identified associations between *Dolosigranulum* and other nasal-associated genera, such as *Corynebacterium*, e.g., [28, 29, 36, 38, 40, 43, 73, 74]. In most cases, the taxonomic resolution in the aforementioned studies was limited to the genus or higher taxonomic levels. Thus, we sought to achieve finer taxonomic resolution and to determine what species are associated with *D. pigrum*. We identified two nostril datasets with V1-V2/V1-V3 16S rRNA gene sequences. These regions contain sufficient information for species-level taxonomic assignment to short-read 16S rRNA gene sequences from most nasal-associated bacteria [26, 41]. After parsing sequences into species-level phylotypes, we interrogated each dataset using Analysis of Composition of Microbiomes (ANCOM) [75] to identify bacterial species that display differential relative abundance in the absence or presence of *D. pigrum* sequences (**Figure 1 and Table S1**). ANCOM is a commonly used approach for identifying associations that accounts for the compositional nature of sequencing data [75]. In the nostrils of 99 children ages 6 and 78 months [26], *Corynebacterium pseudodiphtheriticum* exhibited increased differential relative abundance in the presence of *D. pigrum*, i.e., was positively associated with the presence of *D. pigrum*, as was *Moraxella nonliquefaciens* (**Figure 1A**). In the nostrils of 210 adults from the Human Microbiome Project (HMP), three *Corynebacterium* species––*C. accolens, C. propinquum, C. pseudodiphtheriticum––*and an unresolved supraspecies of *C. accolens-macginleyi-tuberculostearicum* were positively associated with *D. pigrum* (**Figure 1B, panels ii-v**). Whereas, *S. aureus* was negatively associated with *D. pigrum* (**Figure 1B, panel vi**). Our previous analysis of these adult data show these *Corynebacterium* species are the most common *Corynebacterium* species in adult nostrils [41]. Also, all of these *Corynebacterium species* and *D. pigrum* are negatively associated with *S. aureus* in this cohort [41]. Such associations in compositional microbiota data lead to testable hypotheses about possible direct microbe-microbe interactions.

**Figure 1.**
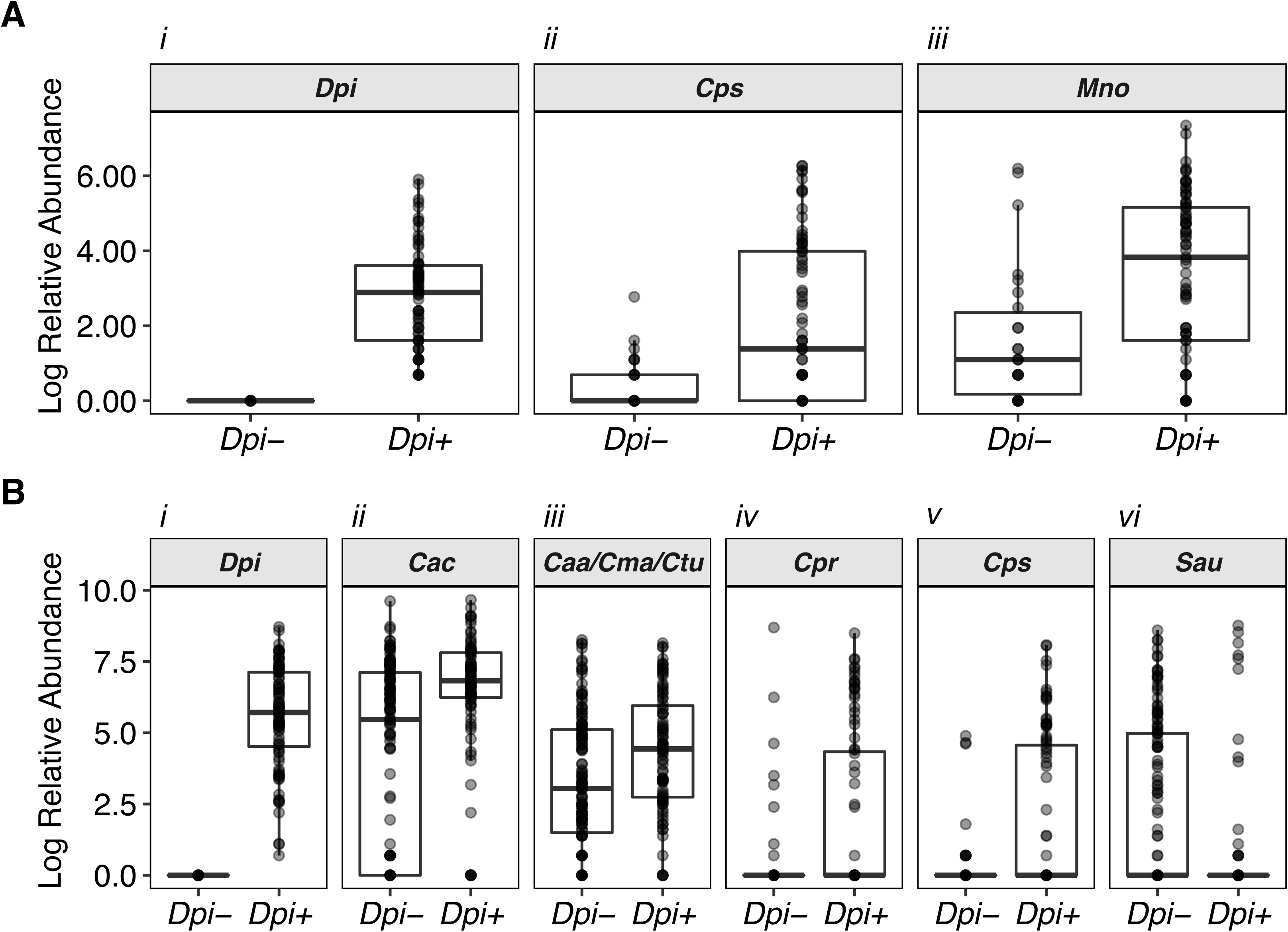
Individual nasal *Corynebacterium* species exhibit increased differential relative abundance in the presence of *D. pigrum* in human nostril microbiota. We used ANCOM to compare species/supraspecies-level composition of 16S rRNA gene nostril datasets from (A) 99 children ages 6 and 78 months and (B) 210 adults when *D. pigrum* was either absent (Dpi-) or present (Dpi+) based on 16S rRNA gene sequencing data. Plots show only the taxa identified as statistically significant (sig = 0.05) after correction for multiple testing within ANCOM. The dark bar represents the median; lower and upper hinges correspond to the first and third quartiles. Each grey dot represents the value for a sample, and multiple overlapping dots appear black. Dpi = *Dolosigranulum pigrum*, Cac = *Corynebacterium accolens*, Caa/Cma/Ctu= supraspecies *Corynebacterium accolens_macginleyi_tuberculostearicum*, Cpr = *Corynebacterium propinquum*, Cps = *Corynebacterium pseudodiphtheriticum*, Mno = *Moraxella nonliquefaciens.* Only three species and one supraspecies of *Corynebacterium* out of the larger number of *Corynebacterium* supraspecies/species present in each dataset met the significance threshold. Specifically, in the adult nostril dataset, there were 21 species and 5 supraspecies groupings of *Corynebacterium* in addition to reads of *Corynebacterium* that were non-assigned (NA) at species level. These data are previously published and visible in Table S7 of reference 42. In the pediatric dataset, there were 16 species of *Corynebacterium* in addition to the (NA) at species level *Corynebacterium* reads (see Table S7 of this manuscript). The Log relative abundance numerical data represented in this figure are available in **Table S1**.

We chose to focus on testing hypotheses about direct interactions between *D. pigrum* and the specific nasal *Corynebacterium* species, as well as between *D. pigrum* and *S. aureus*, for several reasons. First, results from both children and adults identified a positive relationship between *D. pigrum* and individual species of *Corynebacterium* in human nasal microbiota, with a positive association of *D. pigrum* and *C. pseudodiphtheriticum* across age groups (**Figure 1** and **Table S1**). Second, associations between *D. pigrum* and the genus *Corynebacterium* are reported in multiple other human nasal microbiota data sets [28, 29, 36, 38, 40, 43, 73, 74] and, therefore, are more likely to be generalizable and of greater impact for the field. Finally, with respect to possible interactions between *D. pigrum* and *S. aureus,* there is a need to identify potential mechanisms of colonization resistance to *S. aureus* given the lack of an effective vaccine. We then used *in vitro* phenotypic assays to test our hypotheses about direct microbe-microbe interactions.

### Nasal *Corynebacterium* species can enhance the growth of *D. pigrum in vitro*

We hypothesized that the strong positive association between *D. pigrum* and the nasal-associated *Corynebacterium* species might be due to these *Corynebacterium* species releasing metabolites that enhance the growth of *D. pigrum*. As a crude test of this, we quantified *D. pigrum* growth yields on unconditioned agar medium compared to on cell-free agar medium conditioned by growth of *C. pseudodiphtheriticum, C. propinquum* or *C*. *accolens* (**Figure 2**). Conditioning agar medium by prior growth of any of these three nasal *Corynebacterium* species increased the yield (measured as colony forming units, CFUs) of two *D. pigrum* strains (CDC 4709-98 and KPL1914) by one to two orders of magnitude compared to growth on unconditioned agar medium (**Figures 2A** and **2B**). Additionally, one strain of *C. pseudodiphtheriticum* (**Figure 2A**) and the *C. accolens* strain (**Figure 2B**) increased the growth yield of *D. pigrum* CDC 2949-98, a strain with a higher baseline growth yield. The increases in *D. pigrum* growth yield on the *Corynebacterium* cell-free conditioned agar medium could result from either increased growth rate and/or increased viability, and could be consistent with the nasal *Corynebacterium* species either removing a toxin from the medium or releasing a metabolite that enhances growth and/or survival of *D. pigrum*.

**Figure 2.**
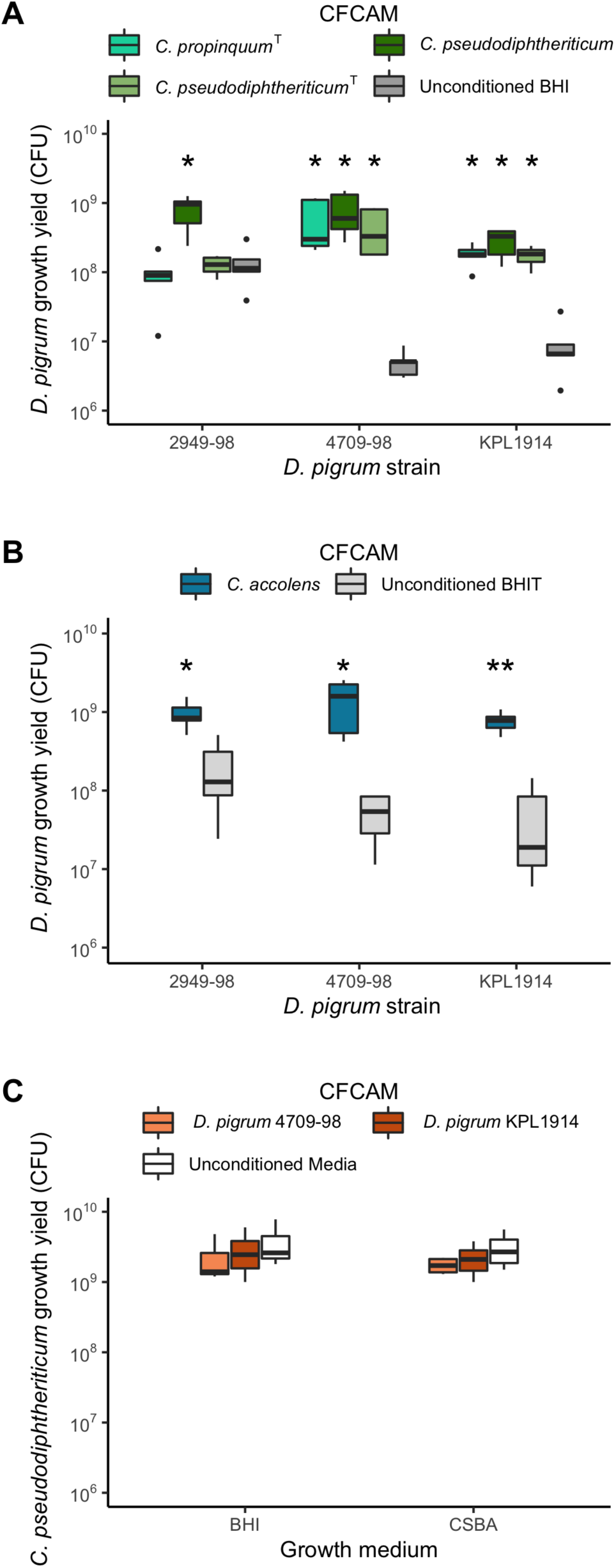
*D. pigrum* growth yields increase on cell-free conditioned agar medium (CFCAM) from nasal *Corynebacterium* species and not vice versa. Growth yield of *D. pigrum* strains CDC 2949-98, CDC 4709-98 and KPL1914 was quantified as the number of CFUs grown on a polycarbonate membrane placed onto (**A**) cell-free conditioned BHI agar from *C. propinquum* (aqua green) or *C. pseudodiphtheriticum* (dark and light green) or (**B**) cell-free conditioned BHI-Triolein (BHIT) agar from *C. accolens* (blue) and compared to growth on unconditioned BHI agar (dark grey) or unconditioned BHIT agar (light grey), respectively. Growth yield of *C. pseudodiphtheriticum* KPL1989 on CFCAM from *D. pigrum* strains (orange) compared to unconditioned medium (white) was assessed similarly (**C**). BHIT was used for growth of *C. accolens* since it is a fatty-acid auxotroph and releases needed oleic acid from triolein. Preconditioning strains were grown on a 0.2-μm, 47-mm polycarbonate membrane for two days to generate CFCAM. After removal, we then placed a new membrane on the CFCAM onto which we spread 100 μL of target bacterial cells that had been resuspended to an OD_600_ of 0.50 in 1x PBS. After 2 days of growth, CFU were enumerated as described in Methods. CFU counts were compared independently for each individual strain (A and B, *n*=5) or medium (C, *n*=4) using a Wilcoxon rank sum test with Bonferroni correction for multiple comparisons to the unconditioned medium. Dark bars represent medians, lower and upper hinges correspond to the first and third quartiles and outlier points are displayed individually. *\**, *p* ≤ 0.05; **, *p* ≤ 0.001

In contrast to the increase in *D. pigrum* growth yield on *C. pseudodiphtheriticum* cell-free conditioned agar medium (**Figure 2A**), there was no increase in *C. pseudodiphtheriticum* strain KPL1989 growth yield on *D. pigrum* cell-free conditioned agar medium (**Figure 2C**). Thus, this growth enhancement goes in one direction from nasal *Corynebacterium* species to *D. pigrum*. This is consistent with unilateral cooperation of nasal *Corynebacterium* species––*C. pseudodiphtheriticum, C. propinquum* or *C. accolens––*with *D. pigrum* in the nostril microbiota and support the observed positive *in vivo* community-level relationships.

The positive association between *C. accolens* and *D. pigrum* in adult nostril microbiota datasets indicates that *in vivo* positive interactions between *C. accolens* and *D. pigrum* prevail (**Figure 1B, panel ii**). However, *in vitro*, we observed either a positive or a negative interaction between *C. accolens* and *D. pigrum* depending on the assay conditions. Unlike *C. propinquum* and *C. pseudodiphtheriticum, C. accolens* is a fatty-acid auxotroph and triolein, a model host epithelial-surface triacylglycerol, serves as a source of needed oleic acid in our assays. We observed increased *D. pigrum* growth yield on a semi-permeable membrane atop *C. accolens* cell-free conditioned agar medium (CFCAM) of Brain Heart Infusion (BHI) supplemented with triolein (BHIT) (**Figure 2B**). In contrast, *D. pigrum* was inhibited when inoculated directly onto this same *C. accolens* cell-free conditioned agar medium (**Table 1**). This inhibition is reminiscent of our previous finding that *in vitro* the *C. accolens*’ triacylglycerol lipase, LipS1, hydrolyzes triacylglycerols releasing free fatty acids that inhibit *S. pneumoniae* [33]. Both *D. pigrum* and *S. pneumoniae* belong to the order Lactobacillales and, based on the closeness of their phylogenetic relationship, we hypothesized that *D. pigrum* might be similarly susceptible to free fatty acids such as the oleic acid that *C. accolens* releases from triolein. Indeed, we observed that oleic acid inhibited *D. pigrum* when we challenged *D. pigrum* with oleic acid using a disk diffusion assay (**Table 2**). We also challenged *D. pigrum* with varying concentrations of oleic acid spread onto plates of BHI agar medium. Similar to the membrane-mediated effect in the *C. accolens* CFCAM experiment above, we observed *D. pigrum* growth at higher concentrations of oleic acid when inoculated onto a semi-permeable membrane atop the oleic-acid-coated medium versus inoculated directly onto the oleic-acid-coated medium (**Table 3**). This indicates the membrane provides some protection from inhibition by oleic acid. Overall, these *in vitro* data indicate that *C. accolens* can both inhibit the growth of *D. pigrum* by releasing antibacterial free fatty acids from host triacylglycerols, such as oleic acid from triolein, (**Tables 1 and 2**) and enhance the growth of *D. pigrum* by releasing an as-yet unidentified factor(s) (**Figure 2B**). Collectively, these results point to a complex set of the molecular interactions between these two species.

**Table 1.** In contrast to when grown on a semi-permeable membrane, *D. pigrum* is inhibited when grown directly on cell-free *C. accolens-*conditioned BHI agar supplemented with triolein as a source of oleic acid.

| Conditioning Strain | Growth of Target Strains<sup>a</sup> |  |  |
| --- | --- | --- | --- |
|  | *S. pneumoniae*<br>603 (6B) | *D. pigrum*<br>CDC 4709-98 | *D. pigrum*<br>KPL1914 |
| *C. accolens*<br>KPL1818 | 0 | 0 | 0 |
| *C. propinquum*<sup>T</sup><br>DSM44285 | 0 | + | + |
| *C. pseudodiphtheriticum*<br>KPL1989 | + | + | + |

<sup>a</sup>0, no growth; +, growth detected, *n*≥3

**Table 2.** Oleic acid inhibits D. pigrum growth.

|  | ZOI (mm)<sup>a</sup> |  |  | 726 |
| --- | --- | --- | --- | --- |
| Oleic Acid<br>(μg/disc) | *S. pneumoniae*<br>603 (6B) | *D. pigrum*<br>CDC 4709-98 | *D. pigrum*<br>KPL1914 |  |
| 20 | 10.3 ± 4.7 | 12.0 ± 2.9 | 17.0 ± 2.1 |  |
| 50 | 22.0 ± 5.4 | 26.8 ± 4.4 | 28.4 ± 7.0 |  |
| 100 | 26.3 ± 6.7 | 35.8 ± 4.5 | 39.4 ± 5.0 |  |

<sup>a</sup>Mean ZOI ± SD produced in a disc-diffusion assay. ZOIs were measured as the smallest diameter of inhibited growth and measurements include disc diameter (6 mm). Biological replicates (*n*=4 for *S. pneumoniae*, *n*=5 for *D. pigrum*) were averaged.

**Table 3.** A 0.2-µm, 47-mm polycarbonate membrane provides *D. pigrum* with some protection against inhibition by oleic acid *in vitro*.

| Oleic acid (μg plated) | Growth directly on agar<sup>a</sup> |  | Growth on membrane |  |
| --- | --- | --- | --- | --- |
|  | *D. pigrum* strain |  |  |  |
|  | KPL1914 | CDC 4709-98 | KPL1914 | CDC 4709-98 |
| 500 | 0 | 0 | 0 | + |
| 50 | 0 | 0 | + | + |
| 5 | + | + | + | + |
| 0.5 | + | + | + | + |
| 0.05 | + | + | + | + |
| 0 (BHI) | + | + | + | + |
| 0 (CSBA) | + | + | + | + |

<sup>a</sup>0, no growth, +, growth detected, *n*=3

### *D*. *pigrum* inhibits *S. aureus* growth

In the absence of a vaccine against *S. aureus*, there are multiple ongoing efforts to identify commensal bacteria that provide colonization resistance to *S. aureus* [15–21, 56] (reviewed in [14]). The ANCOM analysis of the adult nostril microbiota dataset revealed a negative association between *S. aureus* and *D. pigrum* (**Figure 1B, panel vi**), consistent with previous work [31, 41, 56]. Direct antagonism would be the simplest mechanism underpinning this observation. Therefore, we assayed for the effect of 10 different strains of *D. pigrum* on *S. aureus.* We gave *D. pigrum* a head-start to compensate for its slower growth rate *in vitro. S. aureus* growth was inhibited when it was inoculated adjacent to a pregrown inoculum of each of these 10 *D. pigrum* strains on agar medium (**Figure 3**).

**Figure 3.**
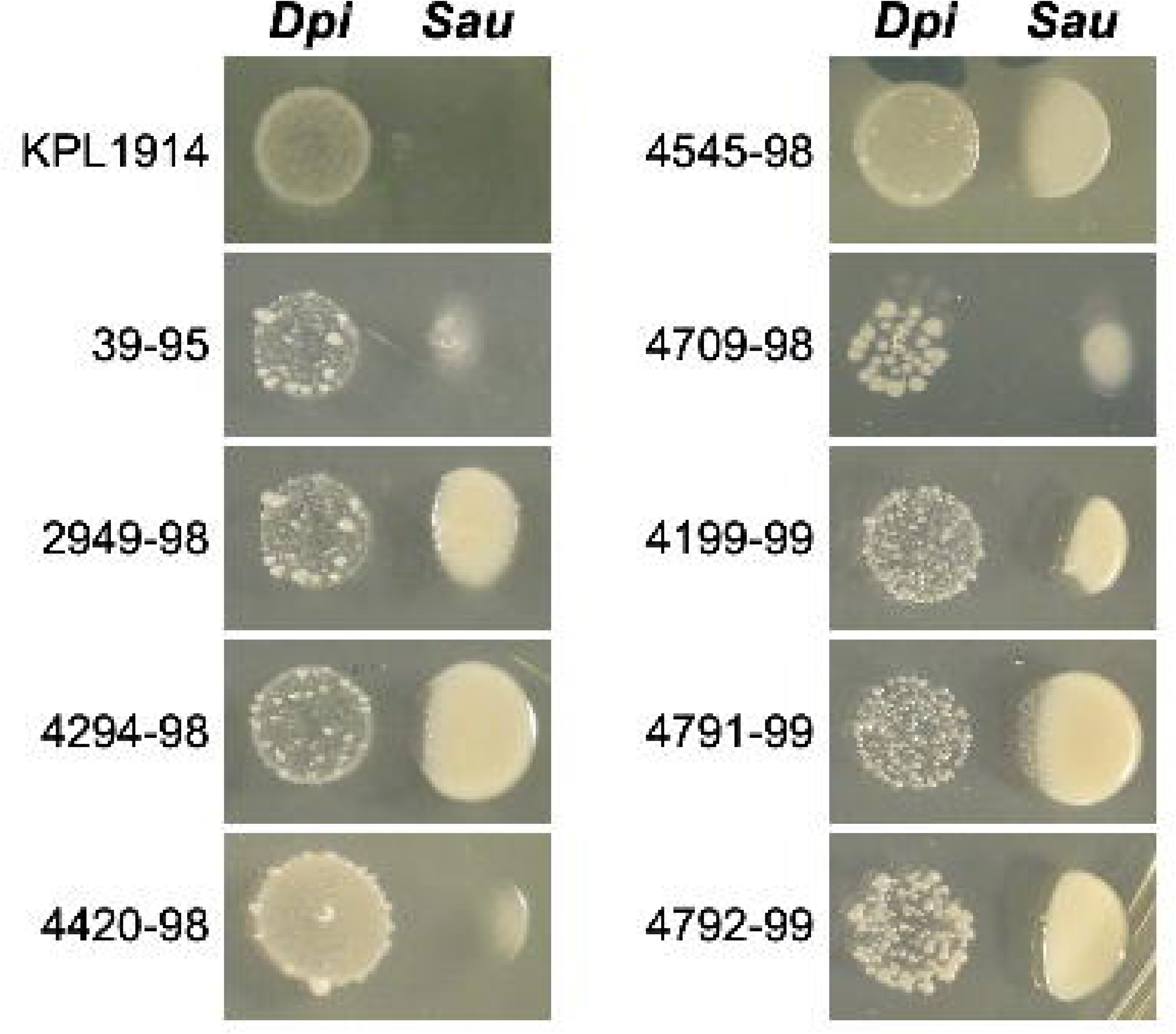
Ten different strains of *D. pigrum* inhibit methicillin-resistant *S. aureus* USA300 strain JE2. Ten pregrown *D. pigrum* isolates produced a diffusible activity that inhibited the growth of *S. aureus* strain JE2 on BHI agar (*n*≥3 independent experiments). Representative images are shown for each strain. *D. pigrum* was resuspended in PBS then a 5 µl drop was placed onto BHI agar and pregrown for 48 hrs. After that, *S. aureus* JE2 was inoculated adjacent to the *D. pigrum*. Inhibition was assessed after 24 and 48 hrs (48 hrs shown here).

### *D. pigrum* production of lactic acid is unlikely to be the primary mechanism for negative associations with *S. pneumoniae* or *S. aureus*

3. *D. pigrum* lactic acid production has been proposed as a mechanism to explain epidemiologic observations of negative associations between *D. pigrum* and *S. pneumoniae* [74]. Under nutrient rich conditions *in vitro*, three tested strains of *D. pigrum* produced from 5.7 to 8.2 mM of L-lactic acid with strain KPL1914 producing the highest concentration (**Figure 4A**). Therefore, we assayed for growth of *S. pneumoniae* in *D. pigrum* KPL1914 cell-free conditioned medium (CFCM) and in BHI broth supplemented with varying concentrations of L-lactic acid. Three of the four *S. pneumoniae* strains tested showed some growth in 22 mM lactic acid (**Figure 4B**), and all strains displayed more growth in BHI supplemented with 11 mM L-lactic acid than in the *D. pigrum* KPL1914 CFCM, which had 7.5 mM of *D. pigrum*-produced L-lactic acid (**Figure 4B**). Thus, the restriction of *S. pneumoniae* growth in *D. pigrum* CFCM is unlikely to be due to *D. pigrum* production of lactic acid. More likely, it reflects competition for nutrients since fresh medium was not added to the CFCM, which, therefore, would have a lower concentration of sugars than BHI broth. However, *D. pigrum* production of a toxin and/or an antipneumococcal compound in BHI broth cannot be excluded. These results indicate that *D. pigrum* production of lactic acid in human nasal passages is unlikely to be the primary molecular mechanism underlying the decreased relative abundance of *S. pneumoniae* in children’s nasal passages when *D. pigrum* is present.

**Figure 4.**
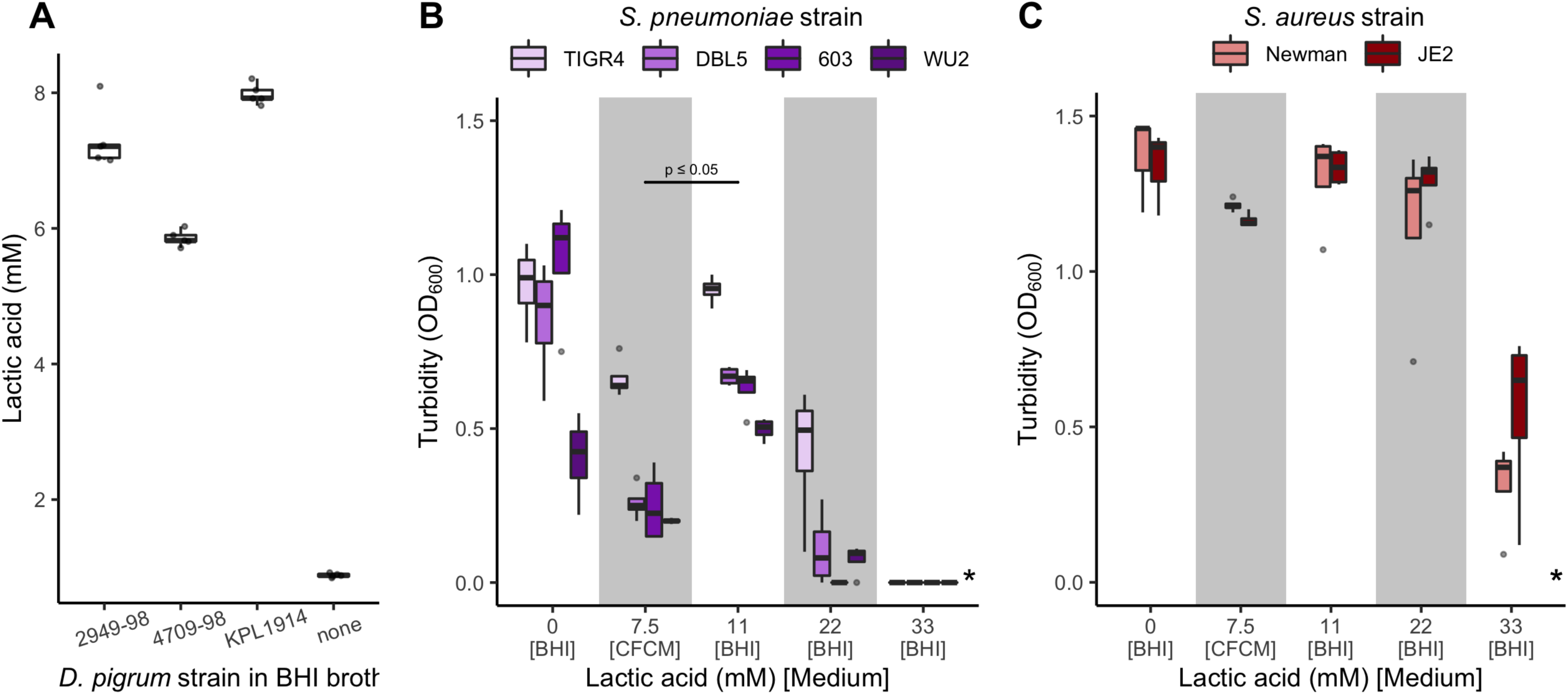
Lactate production by *D. pigrum* is insufficient to inhibit pathobiont growth. Strains of *S. pneumoniae* and *S. aureus* grew in the presence of higher levels of L-lactic acid than those produced by *D. pigrum in vitro*. (**A**) The concentration of L-lactic acid (mM) produced by three *D. pigrum* strains was measured after 24 hrs of gentle shaken aerobic growth in BHI broth at 37°C (*n*=5) as compared to the basal concentration of L-lactic acid in BHI alone (none). (**B**) The average growth (OD_600_) of 4 *S. pneumoniae* strains in *D. pigrum* KPL1914 CFCM or in unconditioned BHI broth supplemented with different concentrations of L-lactic acid measured after 19–20 hrs of static aerobic growth at 37°C (*n*=4). (**C**) The average growth (OD_600_) of 2 *S. aureus* strains in *D. pigrum* KPL1914 CFCM or in unconditioned BHI broth supplemented with different concentrations of L-lactic acid measured after 19–20 hrs of shaken aerobic growth at 37°C (*n*=4). Average growth of *S. pneumoniae* in CFCM and 11 mM L-lactic acid were analyzed independently for each individual strain using a Wilcoxon rank sum test. Dark bars represent medians, lower and upper hinges correspond to the first and third quartiles and outlier points are displayed individually except in panel A where dots for all individual sample values are represented. *None of the *S. pneumoniae* or *S. aureus* strains displayed growth in 55 mM L-lactate.

*D. pigrum* is negatively associated with S. *aureus* in adult nostrils [31, 41, 56] and *D. pigrum* excreted a diffusible activity that inhibited *S. aureus* growth on BHI agar (**Figure 3**). Therefore, we also tested the *in vitro* effect of L-lactic acid on two strains of *S. aureus*. Both showed some growth in 33 mM lactic acid (**Figure 4C**). Thus, under the tested conditions *D. pigrum* does not produce enough L-lactic acid to restrict *S. aureus* growth. In contrast to *S. pneumoniae*, we would not expect depletion of sugars to have a large effect on *S. aureus* growth in *D. pigrum* CFCM given its broader repertoire of energy source utilization options, e.g., amino acids, and indeed both *S. aureus* strains showed little decrease in growth in *D. pigrum* CFCM. This also revealed to a difference in *D. pigrum* production of the anti-*S. aureus* activity during growth on BHI agar medium (**Figure 3**) versus in BHI broth (**Figure 4C**). Excretion of metabolites may vary during growth in liquid versus on agar medium and the mechanism of the *D. pigrum* anti-*S. aureus* activity is yet-to-be identified.

### D. pigrum and C. pseudodiphtheriticum inhibit S. pneumoniae growth together and not alone

Since *C. pseudodiphtheriticum* was positively associated with the presence of *D. pigrum* in both children and adults (**Figure 1**), we investigated the effect of a mixed *in vitro* population of *D. pigrum* and *C. pseudodiphtheriticum* on *S. pneumoniae* growth. Agar medium conditioned with a coculture of *C. pseudodiphtheriticum* strain KPL1989 and *D. pigrum* strain CDC4709-98 inhibited *S. pneumoniae* growth, whereas agar medium conditioned with a monoculture of either *C. pseudodiphtheriticum* or *D. pigrum* alone did not (**Figures 5** and **S1**). This could be due to cocultivation resulting in either a greater level of nutrient competition than monoculture of either commensal alone or in the production of diffusible compound(s) toxic/inhibitory to *S. pneumoniae* by either, or both, *D. pigrum* and/or *C. pseudodiphtheriticum* when grown together. Along with the *Corynebacterium* species enhancement of *D. pigrum* growth yield (**Figure 2**) and the *D. pigrum* inhibition of *S. aureus* growth (**Figure 3**), these data indicate that the negative associations of *D. pigrum* with *S. aureus* and *S. pneumoniae* are likely mediated by different molecular mechanisms.

**Figure 5.**
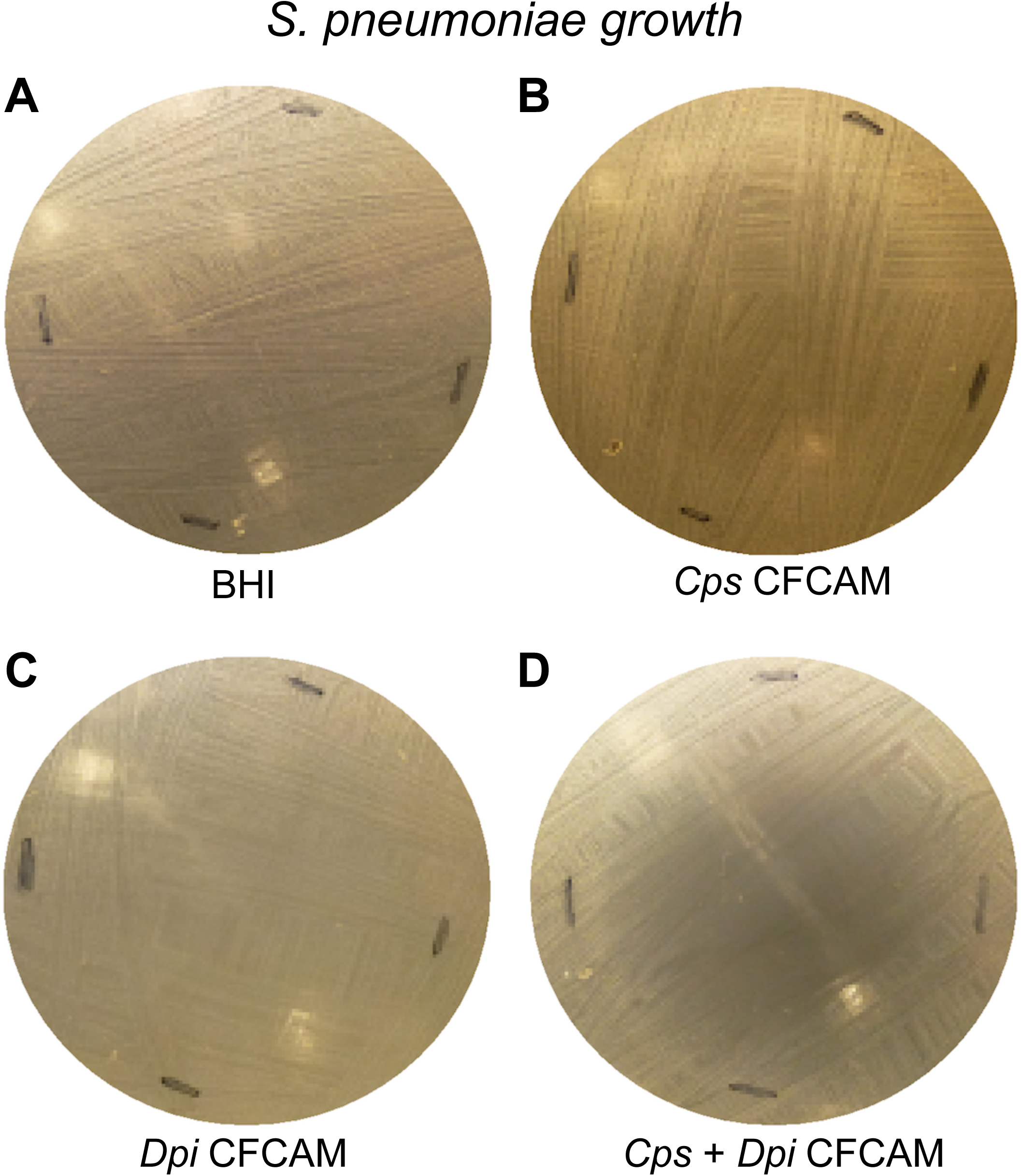
*D. pigrum* and *C. pseudodiphtheriticum* grown together but not *D. pigrum* alone inhibit *S. pneumoniae* in an *in vitro* agar medium-based assay. Representative images of *S. pneumoniae* 603 growth on (**A**) BHI alone or on CFCAM from (**B**) *C. pseudodiphtheriticum* KPL1989, (**C**) *D. pigrum* KPL1914 or (**D**) both *D. pigrum* and *C. pseudodiphtheriticum* grown in a mixed inoculum (*n*=4). To condition the medium, we cultivated *D. pigrum* and/or *C. pseudodiphtheriticum* on a membrane, which was then removed prior to spreading a lawn of *S. pneumoniae*. For monoculture, 100 μL of either *D. pigrum* or *C. pseudodiphtheriticum*, resuspended to an OD_600_=0.50, were inoculated onto the membrane. For mixed coculture, 50 μL of *D. pigrum* (OD_600_=0.50) were mixed with 50 μL of *C. pseudodiphtheriticum* (OD_600_=0.50) to yield a final volume of 100 μL for the inoculum, such that each bacterial species is present in the coculture inoculum at half the amount used for the respective monoculture inoculum. Images were cropped. Black marks indicate edges of where the membrane had been.

Collectively, these phenotypic data (**Figures 2, 3 and 5**) support a role for microbe-microbe interactions in shaping the composition of human nasal microbiota. These also strengthen the case for *D. pigrum* being a beneficial bacterium that can provide colonization resistance against pathobionts. To learn more about the functional capacity and genomic structure of *D. pigrum,* we next turned to genomic analysis, which provided insights into some of the epidemiologic and phenotypic observations presented above.

### The genomes of 11 *D. pigrum* strains reveal a small genome consistent with a highly host-adapted bacterium

We analyzed one publicly available genome of *D. pigrum* (ATCC51524) and sequenced 10 additional strains (**Table S2**), which were selected to ensure representation of distinct strains (see Methods). To start, we focused on basic genomic characteristics. The 11 *D. pigrum* strain genomes had an average size of 1.86 Mb (median 1.88 Mb) with 1693 predicted coding sequences (CDS; **Tables S2** and **S3**). Approximately 1200 CDS were core (**Figures S2 and S3; Table S4**) and exhibited a high degree of nucleotide and amino acid sequence conservation (**Figure S4**). In **Supplemental Text** (**section I**), we further analyzed synteny of two closed genomes (**Figure S5**), did BLAST ring comparisons (**Figure S6**) and constructed a core-gene-based phylogeny (**Figure S7**). The 1.86 Mb genome size is consistent with *D. pigrum* being a highly host-adapted bacterium with reduced biosynthetic capacities, which are detailed below and in the **Supplemental Text** (**section II**) [76].

### *D. pigrum* is a predicted auxotroph for amino acids, polyamines and enzymatic cofactors

The nasal environment is low and/or lacking in key nutrients such as methionine [77] and *D. pigrum*’s small genome size is consistent with reduced biosynthetic capacity. To gain insight into how *D. pigrum* functions in the nasal environment, we examined all 11 genomes finding evidence of auxotrophy for some amino acids (e.g., methionine), polyamines (e.g., putrescine and spermidine) and enzymatic cofactors (e.g., biotin) across all strains. In turn, we identified putative degradation pathways (e.g., methionine), transporters (e.g., polyamines and biotin) and salvage pathways (e.g., folate) suggesting *D. pigrum* acquires some required nutrients exogenously. The **Supplemental Text** (**section II**) contains additional details plus predictions on acquisition of metal cofactors. The auxotrophy predictions may be incomplete, since we were unable to grow *D. pigrum* in a chemically defined medium with all 20 amino acids that was putatively replete based on these predictions. Apparent auxotrophy for a number of required nutrients indicates these must be available either from the host or from neighboring microbes in human nasal passages, e.g., possibly from nasal *Corynebacterium* species.

### Whole genome sequencing indicates that *D. pigrum* metabolizes carbohydrates via homofermentation to lactic acid

*D. pigrum* produced lactate during *in vitro* cultivation (**Figure 4A**). Lactic acid bacteria mainly perform either homo- or heterofermentation of carbohydrates [78]. Therefore, we examined the genomic capacity of *D. pigrum* for carbohydrate metabolism (see also **Supplemental Text, section III**). *D. pigrum* genomes lacked genes required for a complete tricarboxylic acid cycle, which is consistent with fermentation. Moreover, we identified genes encoding a complete glycolytic pathway in all 11 strains that are consistent with homofermentation. All 11 strains harbored a predicted L-lactate-dehydrogenase (EC 1.1.1.27), which catalyzes the reduction of pyruvate to lactate regenerating NAD+ for glycolysis (GAPDH step), consistent with homofermentation to L-lactate as the main product of glycolysis.

### The accessory genome of 11 *D. pigrum* strains contains a diversity of biosynthetic gene clusters predicted to encode antibiotics

Lactic acid production alone appears insufficient to account for the negative *in vitro* associations of *D. pigrum* with *S. aureus* and with *S. pneumoniae* (**Figure 4**). To delve further into the genetic capacity of *D. pigrum* for possible mechanisms of inhibition, we explored the accessory genome of the 11 sequenced strains. Consistent with a prior report [57], *D. pigrum* appears to be broadly susceptible to antibiotics (**Supplement Text, section IV**). What emerged in our analysis was a diversity of biosynthetic gene clusters (BGCs) (**Table S5 and Figure S8**), including a diversity of BGCs predicted to encode candidate antibiotics. Strikingly, although 10 of 10 strains tested displayed inhibition of *S. aureus* growth *in vitro* (**Figure 3**), there was no single BGC common to all 10 strains that might encode a compound with antibiotic activity. Based on this, we hypothesize that *D. pigrum* uses a diverse repertoire of BGCs to produce bioactive molecules that play key roles in interspecies interactions with its microbial neighbors, e.g., for niche competition, and potentially with its host. This points to a new direction for future research on the functions that underlie the positive associations of *D. pigrum* in human nasal microbiota with health and highlights the need to develop a system for genetic engineering of *D. pigrum*.

## Discussion

*D. pigrum* is associated with health in multiple genus-level compositional studies of human URT/nasal passage microbiota. The above species-level genomic and phenotypic experimental data mark a significant advance in the study of *D. pigrum* and set the stage for future research on molecular mechanisms. Further, these phenotypic interactions are consistent with a role for microbe-microbe interactions in shaping the human nasal microbiota. In nasal passage microbiota datasets, we identified positive associations of *D. pigrum* with specific species of *Corynebacterium* in adults and children and a negative association of *D. pigrum* with *S. aureus* in adults (**Figure 1**). We observed phenotypic support for these associations during *in vitro* growth. First, unilateral cooperation from three common nasal *Corynebacterium* species enhanced *D. pigrum* growth yields (**Figure 2**). Second, *D. pigrum* inhibited *S. aureus* (**Figure 3**). Our genomic analysis revealed auxotrophies consistent with *D. pigrum* reliance on cocolonizing microbes and/or the human host for key nutrients. Genomic analysis also showed an aerotolerant anaerobe that performs homofermentation to lactate. However, *D. pigrum* lactate production (**Figure 4A**) was insufficient to inhibit either *S. aureus* (**Figure 4B**) or *S. pneumoniae* (**Figure 4C**), and is, therefore, not the sole contributor to negative associations with *S. pneumoniae* and *S. aureus in vivo*. Consistent with the multiple reports of a negative association between *D. pigrum*, usually in conjunction with the genus *Corynebacterium*, and *S. pneumoniae*, we observed that cocultivation of *D. pigrum* and *C. pseudodiphtheriticum* produced a diffusible activity that robustly inhibited *S. pneumoniae* (**Figures 5** and **S1**) whereas monoculture of either did not. Finally, we uncovered a surprisingly diverse repertoire of BGCs in 11 *D. pigrum* strains, revealing potential mechanisms for niche competition that were previously unrecognized and opening up a new line of investigation in the field.

The *in vitro* interactions of *D. pigrum* with *S. aureus* and with *S. pneumoniae* support inferences from composition-level microbiota data of competition between *D. pigrum* and each pathobiont. However, these interactions differed *in vitro*. *D. pigrum* alone inhibited *S. aureus* but *D. pigrum* plus *C. pseudodiphtheriticum*, together, robustly inhibited *S. pneumoniae*. This points to a more complex set of interactions among these specific bacterial members of the human nasal microbiota, which likely exists in the context of a network of both microbe-microbe and microbe-host interactions. To date, mechanisms for only a few such interactions are described. For example, a *C. accolens* triacylglycerol lipase (LipS1) releases antipneumococcal free fatty acids from model host surface triacyclglycerols *in vitro* pointing to habitat modification as a possible contributor to S*. pneumoniae* colonization resistance [33].

Multiple mechanisms could result in *D. pigrum* inhibition of *S. aureus in vitro* including nutrient competition, excretion of a toxic primary metabolite or of an anti-*S. aureus* secondary metabolite (i.e., an antibiotic). Initial bioassay-guided fractionation approaches failed to identify a mechanism. However, the diverse repertoire of BGCs among the 11 *D. pigrum* strains is intriguing because it includes predicted lanthipeptides and bacteriocins. For example, 4 of the 11 strains harbored putative type II lanthipeptide biosynthetic gene clusters. These clusters are characterized by the presence of the LanM enzyme, containing both dehydration and cyclization domains needed for lanthipeptide biosynthesis [79]. Alignment of these enzymes with the enterococcal cytolysin LanM revealed conserved catalytic residues in both domains [80]. Cleavage of the leader portion of the lanthipeptide is necessary to produce an active compound and the presence of peptidases and transporters within these BGCs suggests these *D. pigrum* strains might secrete an active lanthipeptide, which could play a role in niche competition with other microbes. Additionally, 8 of the 11 *D. pigrum* genomes examined contain putative bacteriocins, or bactericidal proteins and peptides. Intriguingly, the *D. pigrum* strains (CDC4709-98, CDC39-95, KPL1914) exhibiting the strongest inhibition of *S. aureus* (**Figure 3**), were the only strains that contained both a lanthipeptide BGC and a bacteriocin, further indicating that *D. pigrum* may employ multiple mechanisms to inhibit *S. aureus* growth, and if both are required for the *in vitro* inhibition might explain the negative results from bioassay guided fractionation.

Mechanisms are coming to light for how other nasal bacteria interact with *S. aureus*. For example, commensal *Corynebacterium* species excrete a to-be-identified substance that inhibits *S. aureus* autoinducing peptides blocking *agr* quorum sensing (QS) and shifting *S. aureus* shifts towards a commensal phenotype [81]. Also, the to-be-identified mechanism of *C. pseudodiphtheriticum* contact-dependent inhibition of *S. aureus* is mediated through phenol soluble modulins (PSM), the expression of which increases during activation of *agr* QS [82]. Within broader *Staphylococcus-Corynebacterium* interactions, *C. propinquum* outcompetes coagulase-negative *Staphylococcus* (CoNS), but not *S. aureus*, for iron *in vitro* using the siderophore dehydroxynocardamine, the genes for which are transcribed *in vivo* in human nostrils [83]. Interphylum Actinobacteria-Firmicutes interactions also occur between *Cutibacterium acnes* and *Staphylococcus* species (reviewed in [14]). For example, some strains of *C. acnes* produce an anti-staphylococcal thiopeptide, cutimycin, *in vivo* and the presence of the cutimycin BGC is correlated with microbiota composition at the level of the individual human hair follicle [84]. Of note, Actinobacteria competition with coagulase-negative *Staphylococcus* species could also have network-mediated (indirect) effects on *S. aureus* via the well-known competition among *Staphylococcus* species (reviewed in [85]), which can be mediated by antibiotic production, e.g., [15–17, 19], interference with *S. aureus agr* QS [18, 20, 86, 87] or extracellular protease activity [88], among other means [14]. Further rounding out the emerging complexity of microbe-microbe interactions in nasal microbiota, multiple strains of *Staphylococcus*, particularly *S. epidermidis*, inhibit the *in vitro* growth of other nasal and skin bacteria, including *D. pigrum*, via to-be-identified mechanisms [16]. The above points to a wealth of opportunity to use human nasal microbiota as a model system to learn how bacteria use competition to shape their community.

Direct cooperation could contribute to the observed positive associations between bacterial species in epidemiological microbiome studies. Conditioning medium with any of the three nasal *Corynebacterium* species positively associated with *D. pigrum in vivo* in human nasal microbiota (**Figure 1**) enhanced the growth yield of some *D. pigrum* strains (**Figure 2**). This is possibly by excretion of a limiting nutrient or by removal of a toxic medium component. The genomic predictions of auxotrophy (above and supplemental text) might favor nasal *Corynebacterium* species providing cooperation to *D. pigrum* by excretion of a limiting nutrient. Indeed, mass spectrometry indicates a number of nutrients are limiting in the nose [77].

There were several limitations of our study. First, we analyzed the genomes of 11 strains that were primarily isolated in the setting of disease. It is unclear whether these strains were contaminants or pathogenic contributors [57]. However, *D. pigrum* strains are infrequently associated with disease [58-61, 89-92]. These 11 *D. pigrum* strains encoded only a few potential virulence factors, which is consistent with *D. pigrum* acting primarily as a mutualistic species of humans. Second, the ongoing search for a fully defined chemical medium permissive for *D. pigrum* growth precluded experimental verification of predicted auxotrophies and further investigation of how nasal *Corynebacterium* enhance *D. pigrum* growth yields. Third, the *D. pigrum* anti-*S. aureus* factor has eluded purification and identification efforts with standard chemistry approaches and *D. pigrum* is not yet genetically tractable, limiting genetic approaches to identify it. Fourth, to date, there is no animal model for nasal colonization with *D. pigrum* and *Corynebacterium* species, which stymies directly *in vivo* testing the hypothesis of pathobiont inhibition and points to another area of need within the nasal microbiome field.

## Conclusions

In summary, we validated *in vivo* associations from human bacterial microbiota studies with functional assays that support the hypothesis that *D. pigrum* is a mutualist with respect to its human host, rather than a purely commensal bacterium. Further, these phenotypic interactions support a role for microbe-microbe interactions in shaping the composition of human nasal microbiota, and, thus, the possibility of developing microbe-targeted interventions to reshape community composition. The next step will be to identify the molecular mechanisms of those interactions and to assess their role in the human host. Such work could establish the premise for future studies to investigate the therapeutic potential of *D. pigrum* as a topical nasal probiotic for use in patients with recurrent infections with *S. pneumoniae*, possibly in conjunction with a nasal *Corynebacterium* species, or *S. aureus,* in conjunction with established *S. aureus* decolonization techniques [93].

## Methods

### Species-level reanalysis of a pediatric nostril microbiota dataset

Laufer et al. analyzed nostril swabs collected from 108 children ages 6 to 78 months [26]. Of these, 44% were culture positive for *S. pneumoniae* and 23% were diagnosed with otitis media. 16S rRNA gene V1-V2 sequences were generated using Roche/454 with primers 27F and 338R. We obtained 184,685 sequences from the authors, of which 94% included sequence matching primer 338R and 1% included sequence matching primer 27F. We performed demultiplexing in QIIME [94] (split_libraries.py) filtering reads for those ≥250 bp in length, quality score ≥30 and with barcode type hamming_8. Then, we eliminated sequences from samples for which there was no metadata (n=108 for metadata) leaving 120,963 sequences on which we performed *de novo* chimera removal in QIIME (USEARCH 6.1) [95, 96], yielding 120,274 16S rRNA V1-V2 sequences. We then aligned the 120,274 chimera-cleaned reads in QIIIME (PyNAST) [97], using eHOMDv15.04 [41] as a reference database, and trimmed the reads using “o-trim-uninformative-columns-from-alignment” and “o-smart-trim” scripts [98]. 116,620 reads (97% of the chimera-cleaned) were recovered after the alignment and trimming steps. After these initial cleaning steps, we retained only the 99 samples with more than 250 reads. We analyzed this dataset of 99 samples with a total of 114,909 reads using MED [98] with minimum substantive abundance of an oligotype (-M) equal to 4 and maximum variation allowed in each node (-V) equal to 6 nt, which equals 1.6% of the 379-nucleotide length of the trimmed alignment. Of the 114,909 sequences, 82.8% (95,164) passed the -M and -V filtering and are represented in the MED output. Oligotypes were assigned taxonomy in R with the dada2::assignTaxonomy() function (an implementation of the naïve Bayesian RDP classifier algorithm with a kmer size of 8 and a bootstrap of 100) [99, 100] using the eHOMDv15.1 V1-V3 Training Set (version 1) [41] and a bootstrap of 70. We then collapsed oligotypes within the same species/supraspecies yielding the data shown in **Table S6**.

### Microbiota community comparison (Figure 1)

The pediatric 16S rRNA gene V1-V2 dataset analyzed at species level here (**Table S6**), as well as the HMP adult 16S rRNA gene V1-V3 dataset previously analyzed at species level (Table S7 in [41]) were used as input for the ANCOM analysis, including all identified taxa (i.e., we did not remove taxa with low relative abundance). ANCOM (version 1.1.3) was performed using the presence or absence of *D. pigrum,* based on the 16S rRNA gene sequencing data, as group definer. ANCOM default parameters were used (sig = 0.05, tau = 0.02, theta = 0.1, repeated = FALSE (i.e., Kruskal-Wallis test)) except that we performed a correction for multiple comparisons (multcorr = 2), instead of using the default no correction (multcorr = 3) [75]. The Log relative abundance values for the taxa identified as statistically significant (sig = 0.05) are represented in **Figure 1** and also available in **Table S1**.

### Cultivation from frozen stocks

Bacterial strains (**Tables S2 and S7**) were cultivated as described here unless stated otherwise. Across the various methods, strains were grown at 37°C with 5% CO_2_ unless otherwise noted. *D. pigrum* strains were cultivated from frozen stocks on BBL Columbia Colistin-Nalidixic Acid (CNA) agar with 5% sheep blood (BD Diagnostics) for 2 days. *Corynebacterium* species were cultivated from frozen stocks on BHI agar (*C. pseudodiphtheriticum* and *C. propinquum*) or BHI agar supplemented with 1% Tween80 (*C. accolens*) for 1 day. Resuspensions described below were made by harvesting colonies from agar medium and resuspending in 1X phosphate buffered saline (PBS). Of note, we primarily use agar medium because in our experience *D. pigrum* exhibits more consistent growth on agar medium than in liquid medium. Likewise, growth on a semi-solid surface is likely to better represent growth on nasal surfaces than would growth under the well-mixed conditions of shaking liquid medium.

### Preconditioning growth yield assays (Figure 2)

To assess the growth yield of *D. pigrum* on a polycarbonate membrane atop media conditioned by *Corynebacterium* spp. each. *Corynebacterium* strain was resuspended from growth on agar medium to an optical density at 600 nm (OD_600_) of 0.50 in 1x PBS. Then 100 μL of each resuspension was individually spread onto a 0.2-μm, 47-mm polycarbonate membrane (EMD Millipore, Billerica, MA) atop 20 mL of either BHI agar for *C. pseudodiphtheriticum* and *C*. *propinquum* or BHI agar supplemented with Triolein (BHIT) (CAS # 122-32-7, Acros) spread atop the agar medium, as previously described [33], for *C. accolens*. After 2 days of growth, membranes with *Corynebacterium* cells were removed, leaving CFCAM. On each plate of CFCAM, we placed a new membrane onto which we spread 100 μL of *D. pigrum* cells that had been resuspended to an OD_600_ of 0.50 in 1x PBS. After 2 days, membranes with *D. pigrum* were removed, placed in 3 mL 1x PBS, and vortexed for 1 min. to resuspend cells. Resuspensions were diluted 1:10 six times, dilutions were inoculated onto BBL CNA agar with 5% sheep blood and colony forming units (CFUs) were enumerated after 2-3 days of growth. To assess the growth yield of *Corynebacterium pseudodiphtheriticum* on a polycarbonate membrane atop media conditioned by *D. pigrum* strains KPL1914 and CDC 4709-98 were grown for 2 days as described above. *C. pseudodiphtheriticum* KPL1989 growth yield was then measured as described above.

### Growth of *D. pigrum* directly on BHI agar medium supplemented with triolein and conditioned by growth of nasal *Corynebacterium* species (Table 1)

Onto BHI agar supplemented with 200 U/mL of bovine liver catalase (C40-500MG, Sigma) (BHIC), we spread 50 μL of 100 mg/mL of Triolein (BHICT). We then spread 50 μL of a resuspension (OD_600_ of 0.50) of each *Corynebacterium* strain onto a 0.2-μm, 47-mm polycarbonate membrane placed atop 10 mL of BHICT agar in a 100-mm-by-15-mm petri dish. After 2 days, we removed each membrane with *Corynebacterium* cells leaving CFCAM. Using a sterile cotton swab, we then spread either a lawn of *D. pigrum* (from cells resuspended to an OD_600_ of 0.50 in 1x PBS) or *S. pneumoniae* (taken directly from agar medium) onto the CFCAM. Each lawn then grew for 1-2 days before documenting growth or inhibition of growth with digital photography.

### Oleic acid disc diffusion assay (Table 2)

A lawn of *D. pigrum* or *S. pneumoniae* was spread onto 10 mL of BHIC agar using a sterile cotton swab as described above. Oleic acid (Sigma-Aldrich) was dissolved to a final concentration of 2 mg/mL, 5 mg/mL and 10 mg/mL in ethanol and then we added 10 μl of each to separate, sterile 0.2-μm, 6-mm filter discs (Whatman), with 10 μL of ethanol alone added to a disc as a control. After allowing the solvent to evaporate, filter discs were placed onto the bacterial lawns which were then allowed to grow for 1 day before measuring zones of inhibition and photographing.

### Growth of *D. pigrum* directly on versus atop a membrane on oleic-acid-coated agar medium (Table 3)

Oleic was dissolved in 100% ethanol to a concentration of 5 mg/ml and then further diluted 10-fold 5 times in ethanol. For each dilution, 100 μL was spread on top of a separate plate of BHI agar medium. Next, 10 μL of *D. pigrum* KPL1914 and CDC4709-98 each resuspended to OD_600_ = 0.3 was inoculated both directly on the oleic-acid-coated agar medium and atop of a 0.2-µm, 47-mm polycarbonate membrane (EMD Millipore, Billerica, MA) on the same plate. After 2 days at 37°C, we assessed and photographed the growth. In addition, for each dilution and strain one spot on the membrane was resuspended in PBS to assess CFU counts after serial dilutions and plating on blood agar plates (see above).

### *C*. *pigrum–S. aureus* side-by-side coculture assay (Figure 3)

*D. pigrum* cells were harvested with sterile cotton swabs and resuspended in sterile 1x PBS to a minimal OD_600_ of 0.3 then 5 µl drops were individually inoculated on BHI agar medium and incubated for 2 days. *S. aureus* JE2 was grown overnight on BBL Columbia CNA agar with 5% sheep blood and resuspended in PBS to an OD_600_ of 0.1. Then 5 µl drops of *S. aureus* were inoculated at different distances from the pregrown *D. pigrum*. Inhibition was assessed daily and photographically documented.

### Measurement of L-Lactic Acid Concentration (Figure 4A)

*D. pigrum* cells were grown from frozen stocks as above. Cells were then harvested with a sterile cotton swab, resuspended to an OD_600_ of 0.50 in 1x PBS and inoculated at 1:25 in BHI broth for overnight growth gently shaking (∼50-60 rpm) at 37°C under atmospheric conditions. The overnight culture was then inoculated at 1:25 into fresh BHI broth and grown for 24 hrs at 37°C prior to measuring the lactic acid concentration (mmol/L) using a D-lactic acid/L-lactic acid kit per the manufacturer’s instructions (Cat. no. 11112821035, R-Biopharm AG).

### Growth of *S. aureus* and *S. pneumoniae* in *D. pigrum* cell-free conditioned liquid medium

(CFCM in **Figures 4B** and **4C**) After growth in BHI, as described for L-lactic acid measurement, *D. pigrum* KPL1914 cells were removed with a 0.22-μM sterile filter yielding cell-free conditioned medium (CFCM). *S. aureus* strains Newman and JE2 and *S. pneumoniae* strains TIGR4, DBL5, 603, WU2 were each grown on BBL Columbia CNA agar with 5% sheep blood for 1 day, harvested with a sterile cotton swab, resuspended to an OD_600_ of 0.30 in 1x PBS, inoculated at 1:100 into both *D. pigrum* CFCM and BHI broth and grown for 19-20 hrs at 37°C in shaking (*S. aureus*; 50 rpm) or static (*S. pneumoniae*) culture under atmospheric conditions. Growth yield was quantified as OD_600_ absorbance.

### Growth of *S. aureus* and *S. pneumoniae* in BHI broth supplemented with L-lactic acid

(Lactic Acid in **Figures 4B** and **4C**) Strains of *S. aureus* and *S. pneumoniae* were grown and harvested as described above for inoculation. BHI broth, supplemented with L-lactic acid (CAS no. 79-33-4; Fisher BioReagents) at varying concentrations from 11mM – 55 mM, was sterilized through a 0.22-μM filter. After inoculating each strain separately into BHI broth with L-lactic acid, cultures were grown as described above for growth in CFCM. Growth yield was quantified as OD_600_ absorbance.

### Growth assay for *S. pneumoniae* on BHI agar medium conditioned by mono- vs. coculture of *D. pigrum* and/or *C. pseudodiphtheriticum* (Figures 5 and S1)

*D. pigrum* and *C. pseudodiphtheriticum* strains were grown from freezer stocks as described above. Cells were harvested with sterile cotton swabs and resuspended in sterile PBS to an OD_600nm_ of 0.5. We then spotted 100 µl of 1:1 mixed resuspension on a polycarbonate membrane (see above) on BHI agar medium containing 400U/mL bovine liver catalase. After 2 days of growth, the polycarbonate membrane with *D. pigrum* and/or *C. pseudodiphtheriticum* was removed from each plate leaving CFCAM. *S. pneumoniae* 603 [101] was grown overnight on BBL Columbia CNA agar with 5% sheep blood as described above and, using a sterile cotton swab, a lawn was streaked onto the CFCAM and allowed to grow for 24 hours. Growth/inhibition was assessed daily and photographically recorded. Imaging was difficult due to the transparency of *S. pneumoniae* lawns.

### Selection of strains and preparation of DNA for whole genome sequencing

*D. pigrum* KPL1914 was isolated from the nostril of a healthy adult (above). In addition, we selected 9 of 27 *D. pigrum* strains from a CDC collection [57] using an *rpoB*-based typing system with a preference for strains isolated from the nasal passages and/or from children (**Table S2**). Primers Strepto F MOD (AAACTTGGACCAGAAGAAAT) and R MOD (TGTAGCTTATCATCAACCATGTG) were generated *in silico* by mapping primers Strepto F and R [102] to the *rpoB* sequence of *D. pigrum* ATCC 51524 (genome obtained from NCBI; RefSeq: NZ_AGEF00000000.1) with BLAST [103] and manually correcting misalignments in SnapGene viewer 2.8.2 (GSL Biotech, Chicago, IL). PCR were performed using extracted genomic DNA of *D. pigrum*. PCR conditions were as follows: initial denaturation 95°C for 2 minutes, then 30 cycles of denaturation for 30 seconds at 98°C, annealing at 50°C for 30 seconds, elongation 72°C for minutes and a final extension step at 72°C for 10 minutes. PCR products were cleaned using QIAquick PCR purification kit (Qiagen, Germantown, MD) and sequence determined by Sanger sequencing (Macrogen USA, Boston, MA, USA). In the genomic analysis, we also included the publicly available genome for *D. pigrum* ATCC 51524, which was sequenced by the BROAD institute as part of the HMP (RefSeq NZ_AGEF00000000.1).

*D. pigrum* strains were grown atop membranes for 48 hrs as described above. Cells were harvested with a sterile tip, resuspended in 50 µl of sterile PBS and frozen at −80°C. Genomic DNA was extracted using the Epicentre MasterPure nucleic acid extraction kit (Epicentre, Madison, WI) per the manufacturer’s instructions. We assessed DNA purity using a Nanodrop spectrophotometer (Nanodrop, Wilmington, DE), concentration using Qubit fluorometer (Invitrogen, Carlsbad, CA) and fragment size/quality via agarose gel electrophoresis.

### Whole genome sequencing, read assembly, and annotation (Table S3)

Genomic DNA was sequenced at the Yale Center for Genome Analysis (YCGA), New Haven, CT, on an Illumina MiSeq platform using mated paired-end (2 x 250 bp) technology, assembled using de Bruijn graph algorithms with Velvet [104] with a kmer size of 139 bp and annotated with RAST with FIGfam release 70 [105] and Prokka [106]. In addition, *D. pigrum* strains KPL1914 and CDC#4709-98 [57] were sequenced on a PacBio RS II (Pacific Biosystems, Menlo Park, CA) and sequences were assembled using HGAP version 3.0 [107]. We used an iterative procedure to error correct the PacBio genomes, which involved mapping Illumina reads to the PacBio genomes until there were no differences detected between the Illumina reads and the PacBio assembly [108]. To estimate the degree of assembly errors and missing content that might contribute to the variation in gene content, we compared the Illumina assembly of KPL1914 with the Illumina-corrected PacBio assembly of KPL1914 to estimate the possible divergence [109]. Within Illumina assemblies, we identified 139 (1566 vs. 1705) predicted coding sequences as determined by RAST annotation absent in the assembly received by PacBio sequencing. Genomes were deposited at NCBI (GenBank: NAJJ00000000, NAQW00000000, NAQX00000000, NAQV00000000, NAQU00000000, NAQT00000000, NAQS00000000, NAQR00000000, NAQQ00000000 and NAQP00000000 in BioProjects PRJNA379818 and PRJNA379966).

### Identification of the *D. pigrum* core, shell and cloud genome based on Illumina-sequenced genomes from 11 strains (Figures S2 and S3 and Table S4)

Core proteins from RAST-annotated GenBank-files were determined using the intersection of bidirectional best-hits (BDBH), cluster of orthologous (COG) triangles and Markov Cluster Algorithm (OrthoMCL) clustering algorithms using GET_HOMOLOGUES package version 02012019 on Ubuntu-Linux [110] excluding proteins with more than one copy in an input species (as single-copy proteins are safer orthologues, i.e., using flag t-11). GenBank files derived from RAST annotation (see above) were renamed with KPL strain names except for strain ATCC51524. As an initial control, amino acid fasta files (*.faa) were used for the determination of core proteins. We determined the cloud, shell and core genome of each of the 11 sequenced *D. pigrum* strains using the parse_pangenome_matrix.pl script (./parse_pangenome_matrix.pl -m sample_intersection/pangenome_matrix_t0.tab -s) of the GET_HOMOLOGUES package version 30062017 [110]. Definition of cloud, shell and core genome were based on [111]. In brief, cloud is defined as genes only present in a 1 or 2 genomes (cut-off is defined as the class next to the most populated non-core cluster class). The core genome is composed of clusters present in all 11 strains, soft core contains clusters present in 10 genomes and shell includes clusters present in 3 to 9 genomes. Synteny analysis (**Figure S5**) on BDBH core (with flag t11) was performed using the compare_clusters script (-s) and synteny visualization was done in MAUVE using standard settings [112] after the KPL1914 genome was reverse complemented and both genomes had the origin set at the beginning of *dnaA*.

### Phylogenetic reconstruction, sequence and protein similarities

A monophyletic (clade) core genome phylogenic tree was constructed by including *A. otitis* (closest neighbor based on the Living Tree Project [113]) an outgroup (**Figure S7B**). A phylogenic tree without an outgroup was also constructed similarly (**Figure S7A**). *A. otitis* ATCC 51267 contigs were downloaded from NCBI (NZ_AGXA00000000.1) and annotated using RAST (see above). Predicted core proteins common to *A. otitis* and *D. pigrum* genomes were identified as described above using GET_HOMOLOGUES package. Alignments were done using a loop with Clustal Omega V. 1.2.4 ($ for filename in *.faa; do clustalo -i “$filename” -o clustalo_out/${filename%coral} -v; done) and resulting alignments were concatenated using catfasta2phyml perl script (https://github.com/nylander/catfasta2phyml) $./catfasta2phyml.pl *.faa --verbose > outv.phy. PhyML 3.0 [114] with smart model selection [115] using Akaike information criterion was used for phylogenetic analysis (maximum-likelihood) with 100 regular bootstrap replicates and FigTree (http://tree.bio.ed.ac.uk/software/figtree/) for tree visualization.

BLAST Ring Image Generator (BRIG) was used for visualization of the other sequenced genomes compared to the closed CDC 4709-98 genome (**Figure S6**) [116]. Average amino acid and nucleic acid identity (**Figure S4**) was calculated using GET_HOMOLOGUES package version 30062017 [110]. In brief, a pangenome matrix was generated using the OMCL algorithm (./get_homologues.pl -d dpig_folder -t 0 -M (OMCL)) for homologues identification. Both, ANI and AAI were calculated with all available clusters (t 0). Commands used: Generation of an AA identity matrix: $./get_homologues.pl -d “gbk-files” -A -t 0 -M and CDS identity matrix with the command $./get_homologues.pl -d “gbk files” -a ’CDS’ -A -t 0 -M.

### Biosynthetic gene clusters and antibiotic resistance genes (Table S5 and Figure S8)

AntiSMASH (antibiotics & Secondary Metabolite Analysis SHell) and ClusterFinder [117, 118] were accessed at https://antismash.secondarymetabolites.org/ using default setpoints. Putative antibiotic resistance genes or mutations in genes conferring antibiotic resistance were predicted using Resistance Gene Identifier (RGI) on the Comprehensive Antibiotic Resistance Database (CARD) [119]. Assembly contigs were submitted at RGI (https://card.mcmaster.ca/analyze/rgi) and only perfect and strict hits were allowed. ResFinder version 2.1. (https://cge.cbs.dtu.dk/services/ResFinder/) with 90% threshold for %ID and 60% minimum length [120].

### Statistical analyses

R version 3.6.2 was used for statistical analysis and data visualization. The Wilcoxon rank sum test (equivalent to the Mann-Whitney test) was performed using wilcox.test() with paired = FALSE, alternative = “two.sided”.

## List of abbreviations

ANCOM: Analysis of Composition of Microbiomes
CFUs: Colony Forming Units
CFCAM: Cell-Free Conditioned Agar Medium
CFCM: Cell-Free Conditioned Medium
BHI: Brain Heart Infusion
BHIT: Brain Heart Infusion supplemented with Triolein
BGC: Biosynthetic Gene Cluster

## Declarations

### Ethics approval and consent to participate

We isolated *D. pigrum* KPL1914 and *C. pseudodiphtheriticum* KPL1989 from the nostril of an adult as part of a protocol to study the bacterial microbiota of the nostrils of healthy adults that was initially approved by the Harvard Medical School Committee on Human Studies [121], and subsequently approved by the Forsyth Institute Institutional Review Board.

### Consent for publication

Not applicable

### Availability of data and material

The authors declare that all data that support the findings of this study are available within the paper (and its supplementary information files), from publicly available repositories, i.e. GenBank, or from the corresponding authors upon reasonable request. All computer code used in this work is either referenced (for published tools) in the methods section and custom-made code (i.e., loop) is given in the methods section. Further details are available from the corresponding authors on reasonable request

### Authors’ contributions

Conceptualization: SDB, MMP, KPL. Methodology: SDB, SME, MMH. Investigation: SDB, SME, IFE, YK. Interpretation of data: SDB, SME, MMP, IFE, MH, YK, KPL. Visualization: SDB, SME, IFE. Wrote Original Draft: SDB, SME, KPL. Editing and review: SDB, MMP, SME, IFE, KPL. Supervision: SDB, KPL. Funding Acquisition: SDB, MMP, KPL.

## Acknowledgments

We thank Richard R. Facklam and Lynn Shewmaker for providing strains; Joshua Metlay for providing data; Markus Hilty and Stephany Flores Ramos for manuscript edits; and Markus Hilty, Lindsey Bomar, Srikanth Mairpady Shambat, Annelies Zinkernagel and members of the Lemon Lab for helpful discussions.

## REFERENCES

1. Bogaert D, De Groot R, Hermans PW: Streptococcus pneumoniae colonisation: the key to pneumococcal disease. Lancet Infect Dis 2004, 4:144–154.

2. von Eiff C, Becker K, Machka K, Stammer H, Peters G, Group FtS: Nasal carriage as a source of Staphylococcus aureus bacteremia. N Engl J Med 2001, 344:11–16.

3. Wertheim HF, Vos MC, Ott A, van Belkum A, Voss A, Kluytmans JA, van Keulen PH, Vandenbroucke-Grauls CM, Meester MH, Verbrugh HA: Risk and outcome of nosocomial Staphylococcus aureus bacteraemia in nasal carriers versus non-carriers. Lancet 2004, 364:703–705.

4. Kluytmans J, van Belkum A, Verbrugh H: Nasal carriage of *Staphylococcus aureus*: epidemiology, underlying mechanisms, and associated risks. Clin Microbiol Rev 1997, 10:505–520.

5. Young BC, Wu CH, Gordon NC, Cole K, Price JR, Liu E, Sheppard AE, Perera S, Charlesworth J, Golubchik T, et al: Severe infections emerge from commensal bacteria by adaptive evolution. Elife 2017, 6.

6. Bode LG, Kluytmans JA, Wertheim HF, Bogaers D, Vandenbroucke-Grauls CM, Roosendaal R, Troelstra A, Box AT, Voss A, van der Tweel I, et al: Preventing surgical-site infections in nasal carriers of Staphylococcus aureus. N Engl J Med 2010, 362:9–17.

7. Lexau CA, Lynfield R, Danila R, Pilishvili T, Facklam R, Farley MM, Harrison LH, Schaffner W, Reingold A, Bennett NM, et al: Changing epidemiology of invasive pneumococcal disease among older adults in the era of pediatric pneumococcal conjugate vaccine. JAMA 2005, 294:2043–2051.

8. Wahl B, O’Brien KL, Greenbaum A, Majumder A, Liu L, Chu Y, Luksic I, Nair H, McAllister DA, Campbell H, et al: Burden of Streptococcus pneumoniae and Haemophilus influenzae type b disease in children in the era of conjugate vaccines: global, regional, and national estimates for 2000-15. Lancet Glob Health 2018, 6:e744–e757.

9. Collaborators GBDLRI: Estimates of the global, regional, and national morbidity, mortality, and aetiologies of lower respiratory infections in 195 countries, 1990-2016: a systematic analysis for the Global Burden of Disease Study 2016. Lancet Infect Dis 2018, 18:1191–1210.

10. Tong SY, Davis JS, Eichenberger E, Holland TL, Fowler VG, Jr.: Staphylococcus aureus infections: epidemiology, pathophysiology, clinical manifestations, and management. Clin Microbiol Rev 2015, 28:603–661.

11. Turner NA, Sharma-Kuinkel BK, Maskarinec SA, Eichenberger EM, Shah PP, Carugati M, Holland TL, Fowler VG, Jr.: Methicillin-resistant Staphylococcus aureus: an overview of basic and clinical research. Nat Rev Microbiol 2019, 17:203–218.

12. WHO: Pneumococcal vaccines WHO position paper - 2012 - recommendations. Vaccine 2012, 30:4717–4718.

13. Schulfer A, Blaser MJ: Risks of Antibiotic Exposures Early in Life on the Developing Microbiome. PLoS Pathog 2015, 11:e1004903.

14. Brugger SD, Bomar L, Lemon KP: Commensal-Pathogen Interactions along the Human Nasal Passages. PLoS pathogens 2016, 12:e1005633.

15. Zipperer A, Konnerth MC, Laux C, Berscheid A, Janek D, Weidenmaier C, Burian M, Schilling NA, Slavetinsky C, Marschal M, et al: Human commensals producing a novel antibiotic impair pathogen colonization. Nature 2016, 535:511–516.

16. Janek D, Zipperer A, Kulik A, Krismer B, Peschel A: High Frequency and Diversity of Antimicrobial Activities Produced by Nasal Staphylococcus Strains against Bacterial Competitors. PLoS Pathog 2016, 12:e1005812.

17. Nakatsuji T, Chen TH, Narala S, Chun KA, Two AM, Yun T, Shafiq F, Kotol PF, Bouslimani A, Melnik AV, et al: Antimicrobials from human skin commensal bacteria protect against Staphylococcus aureus and are deficient in atopic dermatitis. Sci Transl Med 2017, 9.

18. Paharik AE, Parlet CP, Chung N, Todd DA, Rodriguez EI, Van Dyke MJ, Cech NB, Horswill AR: Coagulase-Negative Staphylococcal Strain Prevents Staphylococcus aureus Colonization and Skin Infection by Blocking Quorum Sensing. Cell Host Microbe 2017, 22:746–756 e745.

19. O’Sullivan JN, Rea MC, O’Connor PM, Hill C, Ross RP: Human skin microbiota is a rich source of bacteriocin-producing staphylococci that kill human pathogens. FEMS Microbiol Ecol 2019, 95.

20. Williams MR, Costa SK, Zaramela LS, Khalil S, Todd DA, Winter HL, Sanford JA, O’Neill AM, Liggins MC, Nakatsuji T, et al: Quorum sensing between bacterial species on the skin protects against epidermal injury in atopic dermatitis. Sci Transl Med 2019, 11.

21. Piewngam P, Zheng Y, Nguyen TH, Dickey SW, Joo HS, Villaruz AE, Glose KA, Fisher EL, Hunt RL, Li B, et al: Pathogen elimination by probiotic Bacillus via signalling interference. Nature 2018, 562:532–537.

22. Tano K, Hakansson EG, Holm SE, Hellstrom S: Bacterial interference between pathogens in otitis media and alpha-haemolytic Streptococci analysed in an in vitro model. Acta Otolaryngol 2002, 122:78–85.

23. Coleman A, Cervin A: Probiotics in the treatment of otitis media. The past, the present and the future. Int J Pediatr Otorhinolaryngol 2019, 116:135–140.

24. Cardenas N, Martin V, Arroyo R, Lopez M, Carrera M, Badiola C, Jimenez E, Rodriguez JM: Prevention of Recurrent Acute Otitis Media in Children Through the Use of Lactobacillus salivarius PS7, a Target-Specific Probiotic Strain. Nutrients 2019, 11.

25. Manning J, Dunne EM, Wescombe PA, Hale JD, Mulholland EK, Tagg JR, Robins-Browne RM, Satzke C: Investigation of Streptococcus salivarius-mediated inhibition of pneumococcal adherence to pharyngeal epithelial cells. BMC Microbiol 2016, 16:225.

26. Laufer AS, Metlay JP, Gent JF, Fennie KP, Kong Y, Pettigrew MM: Microbial communities of the upper respiratory tract and otitis media in children. MBio 2011, 2:e00245–00210.

27. Pettigrew MM, Laufer AS, Gent JF, Kong Y, Fennie KP, Metlay JP: Upper respiratory tract microbial communities, acute otitis media pathogens, and antibiotic use in healthy and sick children. Appl Environ Microbiol 2012, 78:6262–6270.

28. Biesbroek G, Bosch AA, Wang X, Keijser BJ, Veenhoven RH, Sanders EA, Bogaert D: The impact of breastfeeding on nasopharyngeal microbial communities in infants. American journal of respiratory and critical care medicine 2014, 190:298–308.

29. Biesbroek G, Tsivtsivadze E, Sanders EA, Montijn R, Veenhoven RH, Keijser BJ, Bogaert D: Early respiratory microbiota composition determines bacterial succession patterns and respiratory health in children. American journal of respiratory and critical care medicine 2014, 190:1283–1292.

30. Sakwinska O, Bastic Schmid V, Berger B, Bruttin A, Keitel K, Lepage M, Moine D, Ngom Bru C, Brussow H, Gervaix A: Nasopharyngeal microbiota in healthy children and pneumonia patients. J Clin Microbiol 2014, 52:1590–1594.

31. Liu CM, Price LB, Hungate BA, Abraham AG, Larsen LA, Christensen K, Stegger M, Skov R, Andersen PS: Staphylococcus aureus and the ecology of the nasal microbiome. Sci Adv 2015, 1:e1400216.

32. Bosch A, Levin E, van Houten MA, Hasrat R, Kalkman G, Biesbroek G, de Steenhuijsen Piters WAA, de Groot PCM, Pernet P, Keijser BJF, et al: Development of Upper Respiratory Tract Microbiota in Infancy is Affected by Mode of Delivery. EBioMedicine 2016, 9:336–345.

33. Bomar L, Brugger SD, Yost BH, Davies SS, Lemon KP: Corynebacterium accolens Releases Antipneumococcal Free Fatty Acids from Human Nostril and Skin Surface Triacylglycerols. MBio 2016, 7:e01725–01715.

34. Zhang M, Wang R, Liao Y, Buijs MJ, Li J: Profiling of Oral and Nasal Microbiome in Children With Cleft Palate. Cleft Palate Craniofac J 2016, 53:332–338.

35. Salter SJ, Turner C, Watthanaworawit W, de Goffau MC, Wagner J, Parkhill J, Bentley SD, Goldblatt D, Nosten F, Turner P: A longitudinal study of the infant nasopharyngeal microbiota: The effects of age, illness and antibiotic use in a cohort of South East Asian children. PLoS Negl Trop Dis 2017, 11:e0005975.

36. Bosch A, de Steenhuijsen Piters WAA, van Houten MA, Chu M, Biesbroek G, Kool J, Pernet P, de Groot PCM, Eijkemans MJC, Keijser BJF, et al: Maturation of the Infant Respiratory Microbiota, Environmental Drivers, and Health Consequences. A Prospective Cohort Study. Am J Respir Crit Care Med 2017, 196:1582–1590.

37. Kelly MS, Surette MG, Smieja M, Pernica JM, Rossi L, Luinstra K, Steenhoff AP, Feemster KA, Goldfarb DM, Arscott-Mills T, et al: The Nasopharyngeal Microbiota of Children With Respiratory Infections in Botswana. Pediatr Infect Dis J 2017, 36:e211–e218.

38. Hasegawa K, Linnemann RW, Mansbach JM, Ajami NJ, Espinola JA, Petrosino JF, Piedra PA, Stevenson MD, Sullivan AF, Thompson AD, Camargo CA, Jr.: Nasal Airway Microbiota Profile and Severe Bronchiolitis in Infants: A Case-control Study. Pediatr Infect Dis J 2017, 36:1044–1051.

39. Langevin S, Pichon M, Smith E, Morrison J, Bent Z, Green R, Barker K, Solberg O, Gillet Y, Javouhey E, et al: Early nasopharyngeal microbial signature associated with severe influenza in children: a retrospective pilot study. J Gen Virol 2017, 98:2425–2437.

40. Lappan R, Imbrogno K, Sikazwe C, Anderson D, Mok D, Coates H, Vijayasekaran S, Bumbak P, Blyth CC, Jamieson SE, Peacock CS: A microbiome case-control study of recurrent acute otitis media identified potentially protective bacterial genera. BMC Microbiol 2018, 18:13.

41. Escapa IF, Chen T, Huang Y, Gajare P, Dewhirst FE, Lemon KP: New Insights into Human Nostril Microbiome from the Expanded Human Oral Microbiome Database (eHOMD): a Resource for the Microbiome of the Human Aerodigestive Tract. mSystems 2018, 3.

42. Wen Z, Xie G, Zhou Q, Qiu C, Li J, Hu Q, Dai W, Li D, Zheng Y, Wen F: Distinct Nasopharyngeal and Oropharyngeal Microbiota of Children with Influenza A Virus Compared with Healthy Children. Biomed Res Int 2018, 2018:6362716.

43. Copeland E, Leonard K, Carney R, Kong J, Forer M, Naidoo Y, Oliver BGG, Seymour JR, Woodcock S, Burke CM, Stow NW: Chronic Rhinosinusitis: Potential Role of Microbial Dysbiosis and Recommendations for Sampling Sites. Front Cell Infect Microbiol 2018, 8:57.

44. Toivonen L, Hasegawa K, Waris M, Ajami NJ, Petrosino JF, Camargo CA, Jr., Peltola V: Early nasal microbiota and acute respiratory infections during the first years of life. Thorax 2019, 74:592–599.

45. Camelo-Castillo A, Henares D, Brotons P, Galiana A, Rodriguez JC, Mira A, Munoz-Almagro C: Nasopharyngeal Microbiota in Children With Invasive Pneumococcal Disease: Identification of Bacteria With Potential Disease-Promoting and Protective Effects. Front Microbiol 2019, 10:11.

46. Man WH, Clerc M, de Steenhuijsen Piters WAA, van Houten MA, Chu M, Kool J, Keijser BJF, Sanders EAM, Bogaert D: Loss of Microbial Topography between Oral and Nasopharyngeal Microbiota and Development of Respiratory Infections Early in Life. Am J Respir Crit Care Med 2019.

47. Man WH, van Houten MA, Merelle ME, Vlieger AM, Chu M, Jansen NJG, Sanders EAM, Bogaert D: Bacterial and viral respiratory tract microbiota and host characteristics in children with lower respiratory tract infections: a matched case-control study. Lancet Respir Med 2019, 7:417–426.

48. Gan W, Yang F, Tang Y, Zhou D, Qing D, Hu J, Liu S, Liu F, Meng J: The difference in nasal bacterial microbiome diversity between chronic rhinosinusitis patients with polyps and a control population. Int Forum Allergy Rhinol 2019.

49. Perez-Losada M, Alamri L, Crandall KA, Freishtat RJ: Nasopharyngeal Microbiome Diversity Changes over Time in Children with Asthma. PLoS One 2017, 12:e0170543.

50. de Steenhuijsen Piters WAA, Jochems SP, Mitsi E, Rylance J, Pojar S, Nikolaou E, German EL, Holloway M, Carniel BF, Chu M, et al: Interaction between the nasal microbiota and S. pneumoniae in the context of live-attenuated influenza vaccine. Nat Commun 2019, 10:2981.

51. Krismer B, Weidenmaier C, Zipperer A, Peschel A: The commensal lifestyle of Staphylococcus aureus and its interactions with the nasal microbiota. Nat Rev Microbiol 2017, 15:675–687.

52. Man WH, de Steenhuijsen Piters WA, Bogaert D: The microbiota of the respiratory tract: gatekeeper to respiratory health. Nat Rev Microbiol 2017, 15:259–270.

53. Bomar L, Brugger SD, Lemon KP: Bacterial microbiota of the nasal passages across the span of human life. Curr Opin Microbiol 2018, 41:8–14.

54. Esposito S, Principi N: Impact of nasopharyngeal microbiota on the development of respiratory tract diseases. Eur J Clin Microbiol Infect Dis 2018, 37:1–7.

55. Aguirre M, Morrison D, Cookson BD, Gay FW, Collins MD: Phenotypic and phylogenetic characterization of some Gemella-like organisms from human infections: description of Dolosigranulum pigrum gen. nov., sp. nov. J Appl Bacteriol 1993, 75:608–612.

56. Yan M, Pamp SJ, Fukuyama J, Hwang PH, Cho DY, Holmes S, Relman DA: Nasal microenvironments and interspecific interactions influence nasal microbiota complexity and S. aureus carriage. Cell host & microbe 2013, 14:631–640.

57. Laclaire L, Facklam R: Antimicrobial susceptibility and clinical sources of Dolosigranulum pigrum cultures. Antimicrob Agents Chemother 2000, 44:2001–2003.

58. Hoedemaekers A, Schulin T, Tonk B, Melchers WJ, Sturm PD: Ventilator-associated pneumonia caused by Dolosigranulum pigrum. J Clin Microbiol 2006, 44:3461–3462.

59. Lecuyer H, Audibert J, Bobigny A, Eckert C, Janniere-Nartey C, Buu-Hoi A, Mainardi JL, Podglajen I: Dolosigranulum pigrum causing nosocomial pneumonia and septicemia. J Clin Microbiol 2007, 45:3474–3475.

60. Sampo M, Ghazouani O, Cadiou D, Trichet E, Hoffart L, Drancourt M: Dolosigranulum pigrum keratitis: a three-case series. BMC Ophthalmol 2013, 13:31.

61. Venkateswaran N, Kalsow CM, Hindman HB: Phlyctenular keratoconjunctivitis associated with Dolosigranulum pigrum. Ocul Immunol Inflamm 2014, 22:242–245.

62. Song C, Chorath J, Pak Y, Redjal N: Use of Dipstick Assay and Rapid PCR-DNA Analysis of Nasal Secretions for Diagnosis of Bacterial Sinusitis in Children With Chronic Cough. Allergy Rhinol (Providence) 2019, 10:2152656718821281.

63. Chonmaitree T, Jennings K, Golovko G, Khanipov K, Pimenova M, Patel JA, McCormick DP, Loeffelholz MJ, Fofanov Y: Nasopharyngeal microbiota in infants and changes during viral upper respiratory tract infection and acute otitis media. PLoS One 2017, 12:e0180630.

64. Walker RE, Walker CG, Camargo CA, Jr., Bartley J, Flint D, Thompson JMD, Mitchell EA: Nasal microbial composition and chronic otitis media with effusion: A case-control study. PLoS One 2019, 14:e0212473.

65. De Boeck I, Wittouck S, Wuyts S, Oerlemans EFM, van den Broek MFL, Vandenheuvel D, Vanderveken O, Lebeer S: Comparing the Healthy Nose and Nasopharynx Microbiota Reveals Continuity As Well As Niche-Specificity. Front Microbiol 2017, 8:2372.

66. Bogaert D, Keijser B, Huse S, Rossen J, Veenhoven R, van Gils E, Bruin J, Montijn R, Bonten M, Sanders E: Variability and diversity of nasopharyngeal microbiota in children: a metagenomic analysis. PLoS One 2011, 6:e17035.

67. Camarinha-Silva A, Wos-Oxley ML, Jauregui R, Becker K, Pieper DH: Validating T-RFLP as a sensitive and high-throughput approach to assess bacterial diversity patterns in human anterior nares. FEMS Microbiol Ecol 2012, 79:98–108.

68. Wos-Oxley ML, Plumeier I, von Eiff C, Taudien S, Platzer M, Vilchez-Vargas R, Becker K, Pieper DH: A poke into the diversity and associations within human anterior nare microbial communities. ISME J 2010, 4:839–851.

69. Camarinha-Silva A, Jauregui R, Pieper DH, Wos-Oxley ML: The temporal dynamics of bacterial communities across human anterior nares. Environ Microbiol Rep 2012, 4:126–132.

70. Caputo M, Zoch-Lesniak B, Karch A, Vital M, Meyer F, Klawonn F, Baillot A, Pieper DH, Mikolajczyk RT: Bacterial community structure and effects of picornavirus infection on the anterior nares microbiome in early childhood. BMC Microbiol 2019, 19:1.

71. Luna PN, Hasegawa K, Ajami NJ, Espinola JA, Henke DM, Petrosino JF, Piedra PA, Sullivan AF, Camargo CA, Jr., Shaw CA, Mansbach JM: The association between anterior nares and nasopharyngeal microbiota in infants hospitalized for bronchiolitis. Microbiome 2018, 6:2.

72. Peterson SW, Knox NC, Golding GR, Tyler SD, Tyler AD, Mabon P, Embree JE, Fleming F, Fanella S, Van Domselaar G, et al: A Study of the Infant Nasal Microbiome Development over the First Year of Life and in Relation to Their Primary Adult Caregivers Using cpn60 Universal Target (UT) as a Phylogenetic Marker. PLoS One 2016, 11:e0152493.

73. Teo SM, Mok D, Pham K, Kusel M, Serralha M, Troy N, Holt BJ, Hales BJ, Walker ML, Hollams E, et al: The infant nasopharyngeal microbiome impacts severity of lower respiratory infection and risk of asthma development. Cell Host Microbe 2015, 17:704–715.

74. de Steenhuijsen Piters WA, Bogaert D: Unraveling the Molecular Mechanisms Underlying the Nasopharyngeal Bacterial Community Structure. MBio 2016, 7.

75. Mandal S, Van Treuren W, White RA, Eggesbo M, Knight R, Peddada SD: Analysis of composition of microbiomes: a novel method for studying microbial composition. Microb Ecol Health Dis 2015, 26:27663.

76. Pfeiler EA, Klaenhammer TR: The genomics of lactic acid bacteria. Trends Microbiol 2007, 15:546–553.

77. Krismer B, Liebeke M, Janek D, Nega M, Rautenberg M, Hornig G, Unger C, Weidenmaier C, Lalk M, Peschel A: Nutrient limitation governs Staphylococcus aureus metabolism and niche adaptation in the human nose. PLoS Pathog 2014, 10:e1003862.

78. Kandler O: Carbohydrate metabolism in lactic acid bacteria. Antonie Van Leeuwenhoek 1983, 49:209–224.

79. Repka LM, Chekan JR, Nair SK, van der Donk WA: Mechanistic Understanding of Lanthipeptide Biosynthetic Enzymes. Chem Rev 2017, 117:5457–5520.

80. Dong SH, Tang W, Lukk T, Yu Y, Nair SK, van der Donk WA: The enterococcal cytolysin synthetase has an unanticipated lipid kinase fold. Elife 2015, 4.

81. Ramsey MM, Freire MO, Gabrilska RA, Rumbaugh KP, Lemon KP: Staphylococcus aureus Shifts toward Commensalism in Response to Corynebacterium Species. Frontiers in microbiology 2016, 7:1230.

82. Hardy BL, Dickey SW, Plaut RD, Riggins DP, Stibitz S, Otto M, Merrell DS: Corynebacterium pseudodiphtheriticum Exploits Staphylococcus aureus Virulence Components in a Novel Polymicrobial Defense Strategy. MBio 2019, 10.

83. Stubbendieck RM, May DS, Chevrette MG, Temkin MI, Wendt-Pienkowski E, Cagnazzo J, Carlson CM, Gern JE, Currie CR: Competition among Nasal Bacteria Suggests a Role for Siderophore-Mediated Interactions in Shaping the Human Nasal Microbiota. Appl Environ Microbiol 2019, 85.

84. Claesen J, Spagnolo J, Flores Ramos S, Kurita K, Byrd A, Aksenov lA, Melnik A, Wong W, Wang S, Hernandez R, et al: Cutibacterium acnes antibiotic production shapes niche competition in the human skin microbiome. bioRxiv 2019, https://www.biorxiv.org/content/10.1101/594010v1.

85. Parlet CP, Brown MM, Horswill AR: Commensal Staphylococci Influence Staphylococcus aureus Skin Colonization and Disease. Trends Microbiol 2019, 27:497–507.

86. Otto M, Sussmuth R, Vuong C, Jung G, Gotz F: Inhibition of virulence factor expression in Staphylococcus aureus by the Staphylococcus epidermidis agr pheromone and derivatives. FEBS Lett 1999, 450:257–262.

87. Otto M, Echner H, Voelter W, Gotz F: Pheromone cross-inhibition between Staphylococcus aureus and Staphylococcus epidermidis. Infect Immun 2001, 69:1957–1960.

88. Iwase T, Uehara Y, Shinji H, Tajima A, Seo H, Takada K, Agata T, Mizunoe Y: Staphylococcus epidermidis Esp inhibits Staphylococcus aureus biofilm formation and nasal colonization. Nature 2010, 465:346–349.

89. Haas W, Gearinger LS, Hesje CK, Sanfilippo CM, Morris TW: Microbiological etiology and susceptibility of bacterial conjunctivitis isolates from clinical trials with ophthalmic, twice-daily besifloxacin. Adv Ther 2012, 29:442–455.

90. Hall GS, Gordon S, Schroeder S, Smith K, Anthony K, Procop GW: Case of synovitis potentially caused by Dolosigranulum pigrum. J Clin Microbiol 2001, 39:1202–1203.

91. Johnsen BO, Ronning EJ, Onken A, Figved W, Jenum PA: Dolosigranulum pigrum causing biomaterial-associated arthritis. APMIS 2011, 119:85–87.

92. Lin JC, Hou SJ, Huang LU, Sun JR, Chang WK, Lu JJ: Acute cholecystitis accompanied by acute pancreatitis potentially caused by Dolosigranulum pigrum. J Clin Microbiol 2006, 44:2298–2299.

93. Fritz SA, Hogan PG, Hayek G, Eisenstein KA, Rodriguez M, Epplin EK, Garbutt J, Fraser VJ: Household versus individual approaches to eradication of community-associated Staphylococcus aureus in children: a randomized trial. Clinical infectious diseases : an official publication of the Infectious Diseases Society of America 2012, 54:743–751.

94. Caporaso JG, Kuczynski J, Stombaugh J, Bittinger K, Bushman FD, Costello EK, Fierer N, Pena AG, Goodrich JK, Gordon JI, et al: QIIME allows analysis of high-throughput community sequencing data. Nat Methods 2010, 7:335–336.

95. Edgar RC, Haas BJ, Clemente JC, Quince C, Knight R: UCHIME improves sensitivity and speed of chimera detection. Bioinformatics 2011, 27:2194–2200.

96. Edgar RC: Search and clustering orders of magnitude faster than BLAST. Bioinformatics 2010, 26:2460–2461.

97. Caporaso JG, Bittinger K, Bushman FD, DeSantis TZ, Andersen GL, Knight R: PyNAST: a flexible tool for aligning sequences to a template alignment. Bioinformatics 2010, 26:266–267.

98. Eren AM, Morrison HG, Lescault PJ, Reveillaud J, Vineis JH, Sogin ML: Minimum entropy decomposition: unsupervised oligotyping for sensitive partitioning of high-throughput marker gene sequences. The ISME journal 2015, 9:968–979.

99. Callahan BJ, McMurdie PJ, Rosen MJ, Han AW, Johnson AJ, Holmes SP: DADA2: High-resolution sample inference from Illumina amplicon data. Nat Methods 2016, 13:581–583.

100. Wang Q, Garrity GM, Tiedje JM, Cole JR: Naive Bayesian classifier for rapid assignment of rRNA sequences into the new bacterial taxonomy. Appl Environ Microbiol 2007, 73:5261–5267.

101. Malley R, Lipsitch M, Stack A, Saladino R, Fleisher G, Pelton S, Thompson C, Briles D, Anderson P: Intranasal immunization with killed unencapsulated whole cells prevents colonization and invasive disease by capsulated pneumococci. Infect Immun 2001, 69:4870–4873.

102. Drancourt M, Roux V, Fournier PE, Raoult D: rpoB gene sequence-based identification of aerobic Gram-positive cocci of the genera Streptococcus, Enterococcus, Gemella, Abiotrophia, and Granulicatella. J Clin Microbiol 2004, 42:497–504.

103. Altschul SF, Gish W, Miller W, Myers EW, Lipman DJ: Basic local alignment search tool. J Mol Biol 1990, 215:403–410.

104. Zerbino DR, Birney E: Velvet: algorithms for de novo short read assembly using de Bruijn graphs. Genome Res 2008, 18:821–829.

105. Aziz RK, Bartels D, Best AA, DeJongh M, Disz T, Edwards RA, Formsma K, Gerdes S, Glass EM, Kubal M, et al: The RAST Server: rapid annotations using subsystems technology. BMC Genomics 2008, 9:75.

106. Seemann T: Prokka: rapid prokaryotic genome annotation. Bioinformatics 2014, 30:2068–2069.

107. Chin CS, Alexander DH, Marks P, Klammer AA, Drake J, Heiner C, Clum A, Copeland A, Huddleston J, Eichler EE, et al: Nonhybrid, finished microbial genome assemblies from long-read SMRT sequencing data. Nat Methods 2013, 10:563–569.

108. Pettigrew MM, Ahearn CP, Gent JF, Kong Y, Gallo MC, Munro JB, D’Mello A, Sethi S, Tettelin H, Murphy TF: Haemophilus influenzae genome evolution during persistence in the human airways in chronic obstructive pulmonary disease. Proc Natl Acad Sci U S A 2018, 115:E3256–E3265.

109. Hilty M, Wuthrich D, Salter SJ, Engel H, Campbell S, Sa-Leao R, de Lencastre H, Hermans P, Sadowy E, Turner P, et al: Global phylogenomic analysis of nonencapsulated Streptococcus pneumoniae reveals a deep-branching classic lineage that is distinct from multiple sporadic lineages. Genome Biol Evol 2014, 6:3281–3294.

110. Contreras-Moreira B, Vinuesa P: GET_HOMOLOGUES, a versatile software package for scalable and robust microbial pangenome analysis. Appl Environ Microbiol 2013, 79:7696–7701.

111. Kaas RS, Friis C, Ussery DW, Aarestrup FM: Estimating variation within the genes and inferring the phylogeny of 186 sequenced diverse Escherichia coli genomes. BMC Genomics 2012, 13:577.

112. Darling AE, Mau B, Perna NT: progressiveMauve: multiple genome alignment with gene gain, loss and rearrangement. PLoS One 2010, 5:e11147.

113. Yarza P, Richter M, Peplies J, Euzeby J, Amann R, Schleifer KH, Ludwig W, Glockner FO, Rossello-Mora R: The All-Species Living Tree project: a 16S rRNA-based phylogenetic tree of all sequenced type strains. Syst Appl Microbiol 2008, 31:241–250.

114. Guindon S, Dufayard JF, Lefort V, Anisimova M, Hordijk W, Gascuel O: New algorithms and methods to estimate maximum-likelihood phylogenies: assessing the performance of PhyML 3.0. Syst Biol 2010, 59:307–321.

115. Lefort V, Longueville JE, Gascuel O: SMS: Smart Model Selection in PhyML. Mol Biol Evol 2017, 34:2422–2424.

116. Alikhan NF, Petty NK, Ben Zakour NL, Beatson SA: BLAST Ring Image Generator (BRIG): simple prokaryote genome comparisons. BMC Genomics 2011, 12:402.

117. Medema MH, Blin K, Cimermancic P, de Jager V, Zakrzewski P, Fischbach MA, Weber T, Takano E, Breitling R: antiSMASH: rapid identification, annotation and analysis of secondary metabolite biosynthesis gene clusters in bacterial and fungal genome sequences. Nucleic acids research 2011, 39:W339–346.

118. Weber T, Blin K, Duddela S, Krug D, Kim HU, Bruccoleri R, Lee SY, Fischbach MA, Muller R, Wohlleben W, et al: antiSMASH 3.0-a comprehensive resource for the genome mining of biosynthetic gene clusters. Nucleic Acids Res 2015, 43:W237–243.

119. McArthur AG, Waglechner N, Nizam F, Yan A, Azad MA, Baylay AJ, Bhullar K, Canova MJ, De Pascale G, Ejim L, et al: The comprehensive antibiotic resistance database. Antimicrob Agents Chemother 2013, 57:3348–3357.

120. Zankari E, Hasman H, Cosentino S, Vestergaard M, Rasmussen S, Lund O, Aarestrup FM, Larsen MV: Identification of acquired antimicrobial resistance genes. J Antimicrob Chemother 2012, 67:2640–2644.

121. Lemon KP, Klepac-Ceraj V, Schiffer HK, Brodie EL, Lynch SV, Kolter R: Comparative analyses of the bacterial microbiota of the human nostril and oropharynx. MBio 2010, 1:e00129–00110

