## SupplementalMaterial for "*Dolosigranulum pigrum* cooperation and competition in human nasal microbiota"

**Table S1A. Adult nostril dataset ANCOM results. These Log abundance values are plotted in Figure 1A.** (See excel file)

**Table S1B. Pediatric nostril dataset ANCOM results. These Log abundance values are plotted in Figure 1B.** (See excel file)

**Figure S1. When grown together *D. pigrum* and *C. pseudodiphtheriticum* inhibit *S. pneumoniae*.** A second set of representative images of *S. pneumoniae* 603 growth on (**A**) BHI alone or CFCAM from (**B**) *C.* *pseudodiphtheriticum* KPL1989, (**C**) *D. pigrum* KPL1914 or (**D**) both *D. pigrum* and *C. pseudodiphtheriticum* grown in a mixed inoculum (*n*=4). To condition the medium, we cultivated *D. pigrum* and/or *C. pseudodiphtheriticum* on a membrane, which was then removed prior to spreading a lawn of *S. pneumoniae*. Images were cropped. Black marks indicate edges of where the membrane had been.

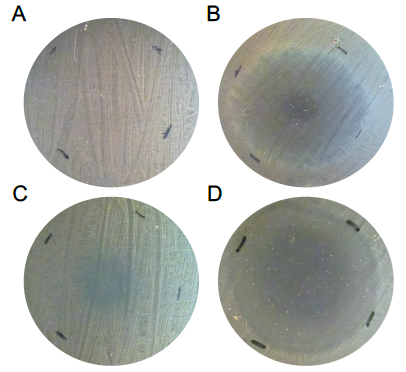

**Table S2. Characteristics of *D. pigrum* strains and Illumina genomes in this study.**

| **Strain name** | **Internal strain #** | **Source (citation)** | **Geography / body site / age** | **Median fold coverage** | **CDS**^+^ | **RNAs**^+^ | **GC (%)**^+^ | **Size (bp)** |
| --- | --- | --- | --- | --- | --- | --- | --- | --- |
| KPL1914 | KPL1914 | This study | Massachusetts / nostril / adult | 83 | 1566 | 72 | 40 | 1,726,398 |
| CDC 39-95 | KPL1922 | CDC [1] | Canada / sinus / 3 years | 62 | 1666 | 80 | 39.7 | 1,859,258 |
| CDC 2949-98 | KPL1930 | CDC [1] | AZ / nasopharyngeal / NA | 60 | 1644 | 73 | 39.6 | 1,886,398 |
| CDC 4294-98 | KPL1931 | CDC [1] | SC / blood / 2 months | 73 | 1841 | 78 | 39.5 | 2,014,679 |
| CDC 4420-98 | KPL1932 | CDC [1] | TN / blood / 11 years | 63 | 1745 | 73 | 39.7 | 1,934,436 |
| CDC 4545-98 | KPL1933 | CDC [1] | AZ / nasopharyngeal / NA | 128 | 1680 | 61 | 39.6 | 1,861,299 |
| CDC 4709-98 | KPL1934 | CDC [1] | GA / eye / 2 months | 81 | 1686 | 80 | 39.6 | 1,912,682 |
| CDC 4199-99 | KPL1937 | CDC [1] | GA / blood / 1.8 years | 107 | 1746 | 85 | 39.6 | 1,976,602 |
| CDC 4791-99 | KPL1938 | CDC [1] | AZ / nasopharyngeal / NA | 61 | 1651 | 80 | 39.6 | 1,873,869 |
| CDC 4792-99 | KPL1939 | CDC [1] | AZ / nasopharyngeal / NA | 92 | 1704 | 80 | 39.4 | 1,893,917 |
| SS-1342 (NCFB2967) (ATCC 51524) | NA | BROAD / HMP R91/1468 [2] | England / spinal cord (autopsy) | 200* | 1651 | 31 | 39.6 | 1,846,028 |

***sequenced by the BROAD institute for the Human Microbiota Project**

**^+^ as determined by RAST annotation (see text)**

**Table S3. Assembly characteristics of *D. pigrum* Illumina genomes in this study.**

| **CDC / internal strain #** | | **Nodes [N]** | **N50 [bp]** | **Max [bp]** | **Total [bp]** | **Reads [N]** |
| --- | --- | --- | --- | --- | --- | --- |
|  | KPL1914 | 107 | 87,900 | 210,575 | 1,726,398 | 1,640,818 |
| 39-95 | KPL1922 | 138 | 153,015 | 273,357 | 1,859,258 | 1,819,350 |
| 2949-98 | KPL1930 | 179 | 127,744 | 456,484 | 1,886,398 | 1,699,318 |
| 4294-98 | KPL1931 | 139 | 88,498 | 198,469 | 2,014,679 | 2,357,776 |
| 4420-98 | KPL1932 | 134 | 209,743 | 328,871 | 1,934,436 | 2,098,746 |
| 4545-98 | KPL1933 | 50 | 283,724 | 492,087 | 1,861,299 | 2,738,030 |
| 4709-98 | KPL1934 | 142 | 110,284 | 268,980 | 1,912,682 | 2,198,156 |
| 4199-99 | KPL1937 | 109 | 128,019 | 379,812 | 1,976,602 | 2,528,652 |
| 4791-99 | KPL1938 | 129 | 132,767 | 316,186 | 1,873,869 | 1,831,854 |
| 4792-99 | KPL1939 | 86 | 253,067 | 460,850 | 1,893,917 | 2,039,094 |

**Figure S2. The conservative core genome of 11 *D. pigrum* strains encodes 1200 orthologs.** A Venn diagram generated using the bidirectional best-hits (BDBH), cluster of orthologous groups (COG) triangle and OrthoMCL (OMCL) algorithms identified predicted protein orthologs (RAST annotation) shared between the 11 *D. pigrum* genomes (GET_HOMOLOGUES package version 02012019) [3]. Flag t=11 was used to only include clusters containing single-copy orthologs from all input species since these are likely the most reliable ortholog predictions.

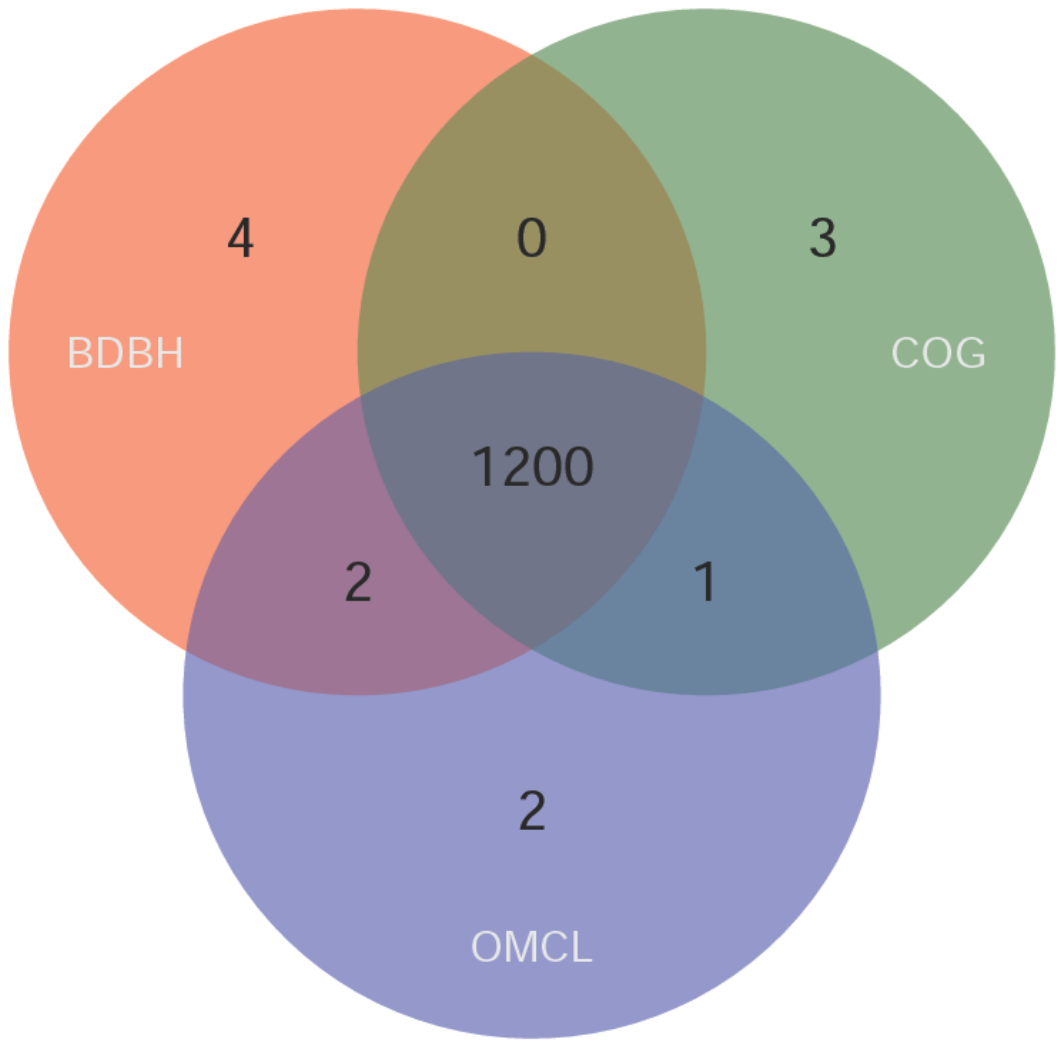

**Figure S3. Core, shell and cloud pangenome of 11 *Dolosigranulum pigrum* strains.** The pangenome of the 11 *D. pigrum* strains includes an estimated core of 1216, soft core of 116, shell of 373 and cloud of 1024 genes (CDS), as determined by the parse_pangenome_matrix.pl script (using the OMCL / COG intersection) of the GET_HOMOLOGUES package version 02012019 [3]. The core genome is composed of genes that are present in all strains and soft core contains clusters present in 10 genomes but not the core as defined in [4]. Cloud is defined as genes only present in a few genomes (cut-off is defined as the class next to the most populated non-core cluster class). Shell genes are the remaining genes and displayed sorted to the number of genomes in which these are present (*n*=3-9).

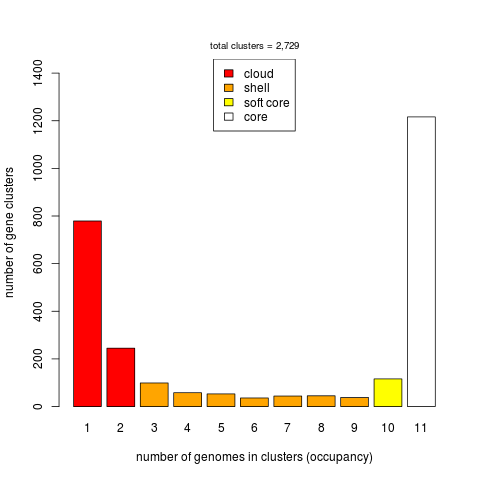

**Table S4. Distribution by strain of genes in the core, soft core, shell and cloud *D. pigrum* genome.***

| **CDC or other strain #** | **Internal reference** | **Cloud [N]** | **Shell [N]** | **Soft core [N]** | **Core [N]** |
| --- | --- | --- | --- | --- | --- |
| KPL1914 | KPL1914 | 79 | 135 | 89 | 1216 |
| 39-95 | KPL1922 | 81 | 189 | 109 | 1216 |
| 2949-98 | KPL1930 | 88 | 167 | 113 | 1216 |
| 4294-98 | KPL1931 | 243 | 183 | 110 | 1216 |
| 4420-98 | KPL1932 | 134 | 205 | 114 | 1216 |
| 4545-98 | KPL1933 | 102 | 202 | 96 | 1216 |
| 4709-98 | KPL1934 | 105 | 185 | 111 | 1216 |
| 4199-99 | KPL1937 | 186 | 174 | 94 | 1216 |
| 4791-99 | KPL1938 | 73 | 192 | 108 | 1216 |
| 4792-99 | KPL1939 | 110 | 194 | 108 | 1216 |
| ATCC51524 |  | 68 | 194 | 108 | 1216 |

* These results were determined as described for Figure S3.

**Figure S4. The (A) average nucleotide identities (ANI) and (B) amino acid identities (AAI) show a high degree of conservation across all 11 *D. pigrum* genomes.** Average identity matrices of clustered coding sequences were calculated using GET_HOMOLOGUES with the OrthoMCL algorithm. Both ANI and AAI were calculated with all available clusters (t 0). Commands used: Generation of an AA identity matrix: $ ./get_homologues.pl -d “gbk-files” -A -t 0 -M and CDS identity matrix with the command $./get_homologues.pl -d “gbk files” -a 'CDS' -A -t 0 -M.

**A**

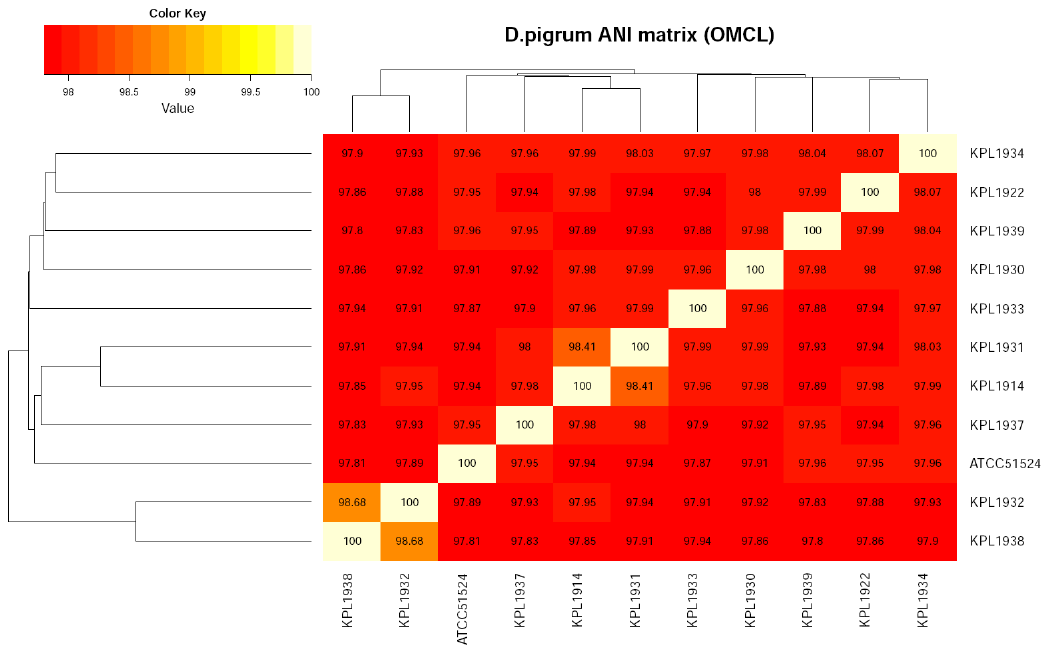

**B**

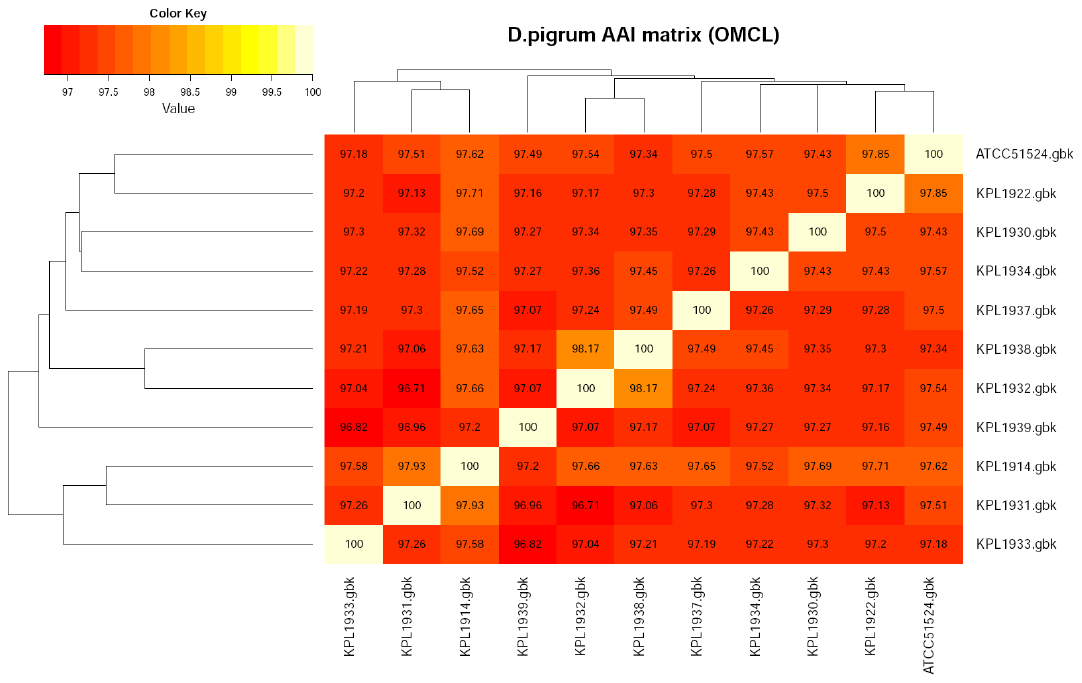

**Figure S5. Comparative analysis of two *D. pigrum* genomes reveals a high degree of synteny.** Multiple genome alignment for synteny analysis of the two closed *D. pigrum* genomes CDC 4709-98 and KPL1914 was performed using MAUVE [5]. Locally collinear blocks (color coded) and similarity profiles are presented, and genome boundaries are indicated (red bar after non-aligned sequences, i.e., sequences that are unique for the corresponding genome).

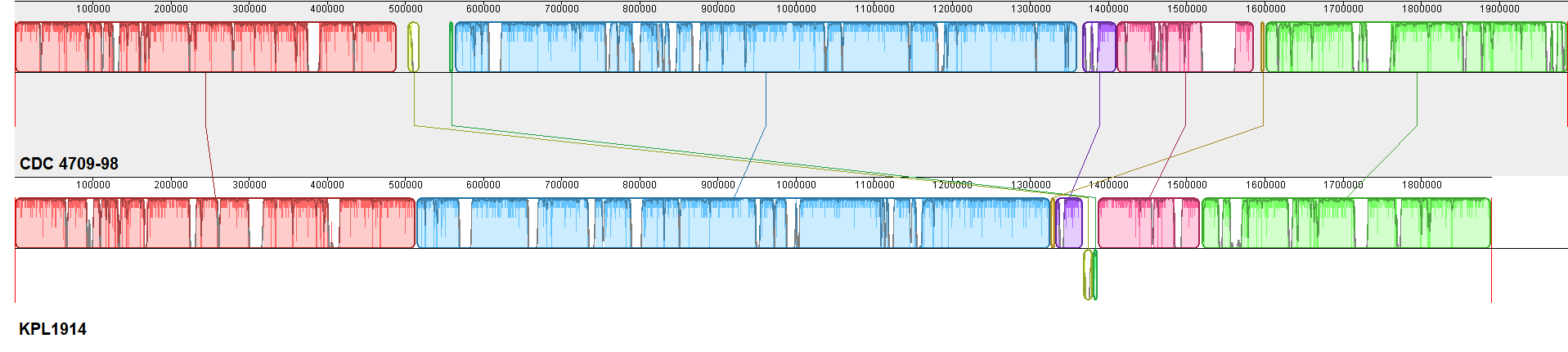

**Figure S6. *D. pigrum* genomes BLAST ring comparisons.** BLAST Ring Image Generator (BRIG) version 0.95 [6] was used for visualization of the sequenced genomes with the closed genome of CDC 4709-98 as a reference. (**A**) Closed genome of CDC 4709-98 with GC plots (**B**) Closed genome of KPL 1914 with GC plots (**C**) BLASTN based comparison of the closed genomes of CDC 4709-98 and KPL 1914 (**D**) BLASTN based comparison of CDC 4709-98 and the remaining 10 Illumina contig-based genomes as well as ATCC 51524.

**5A**

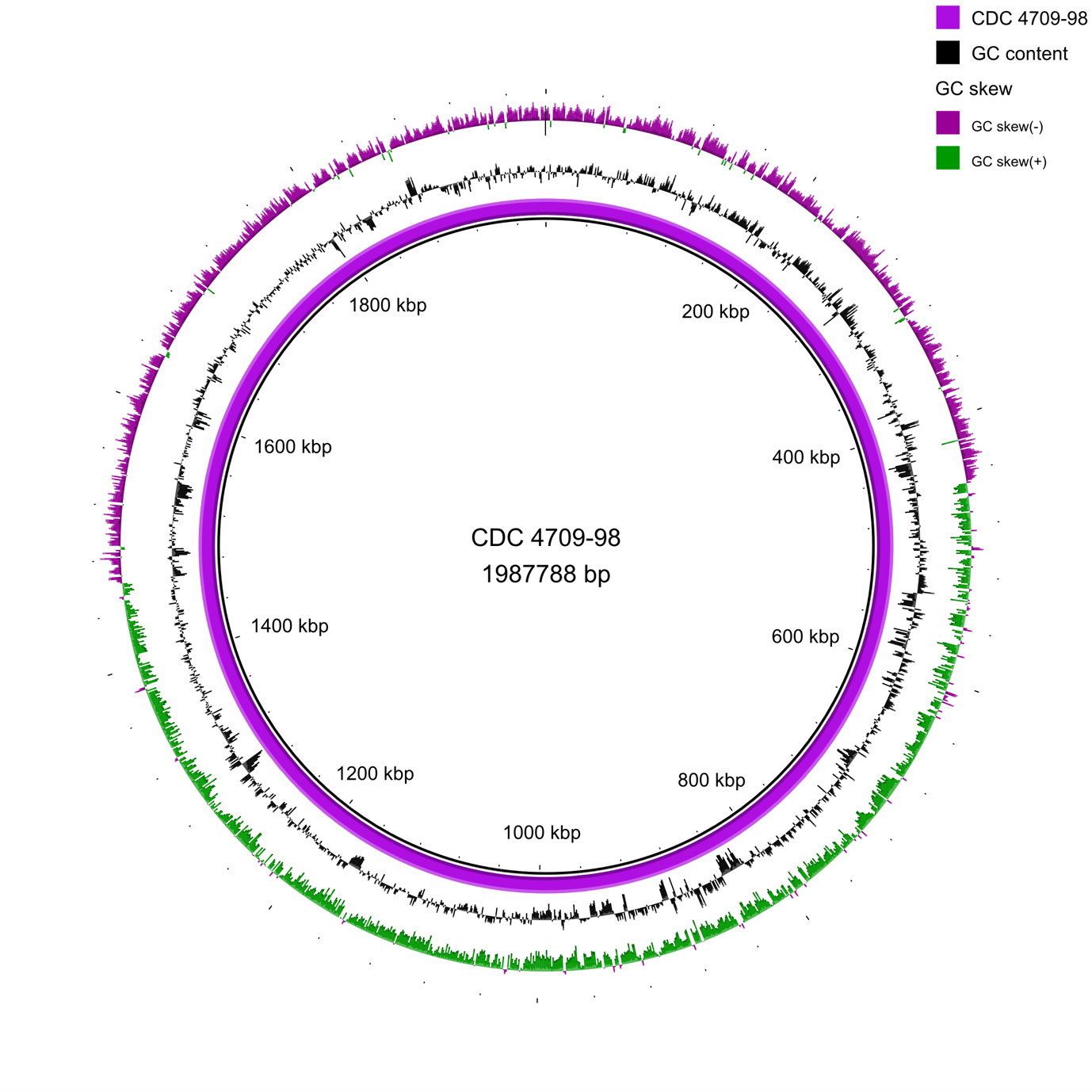

**5B**

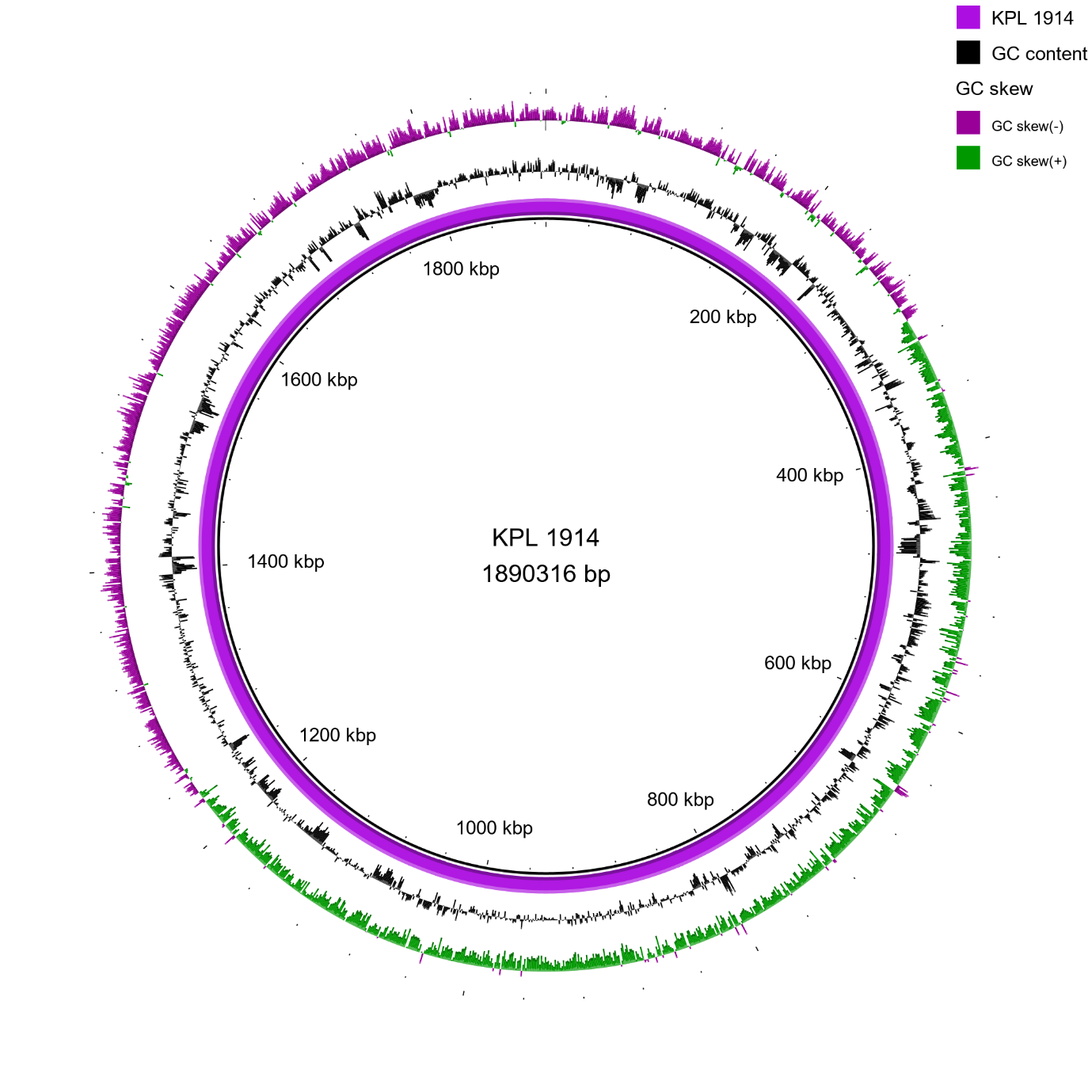

**5C**

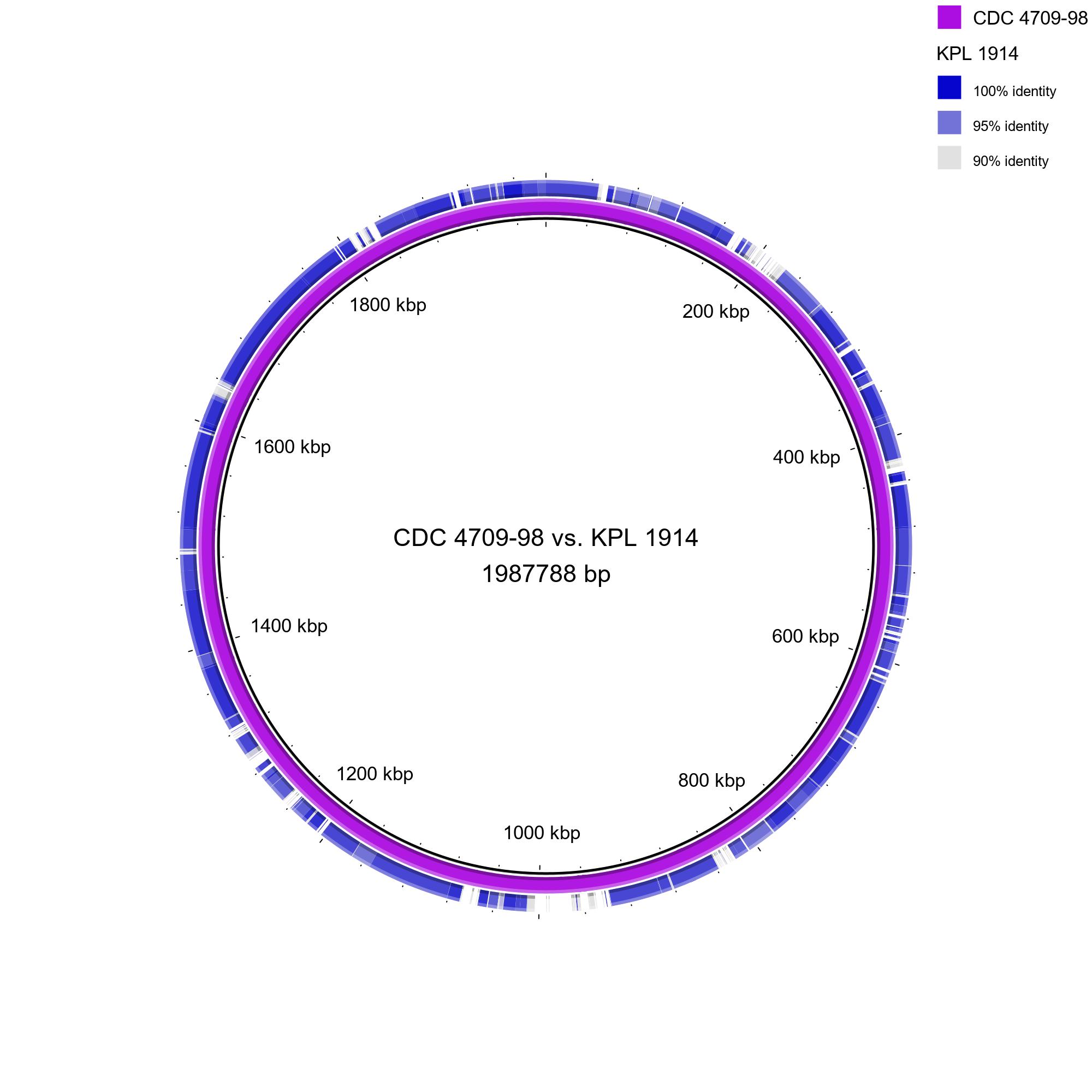

**5D**

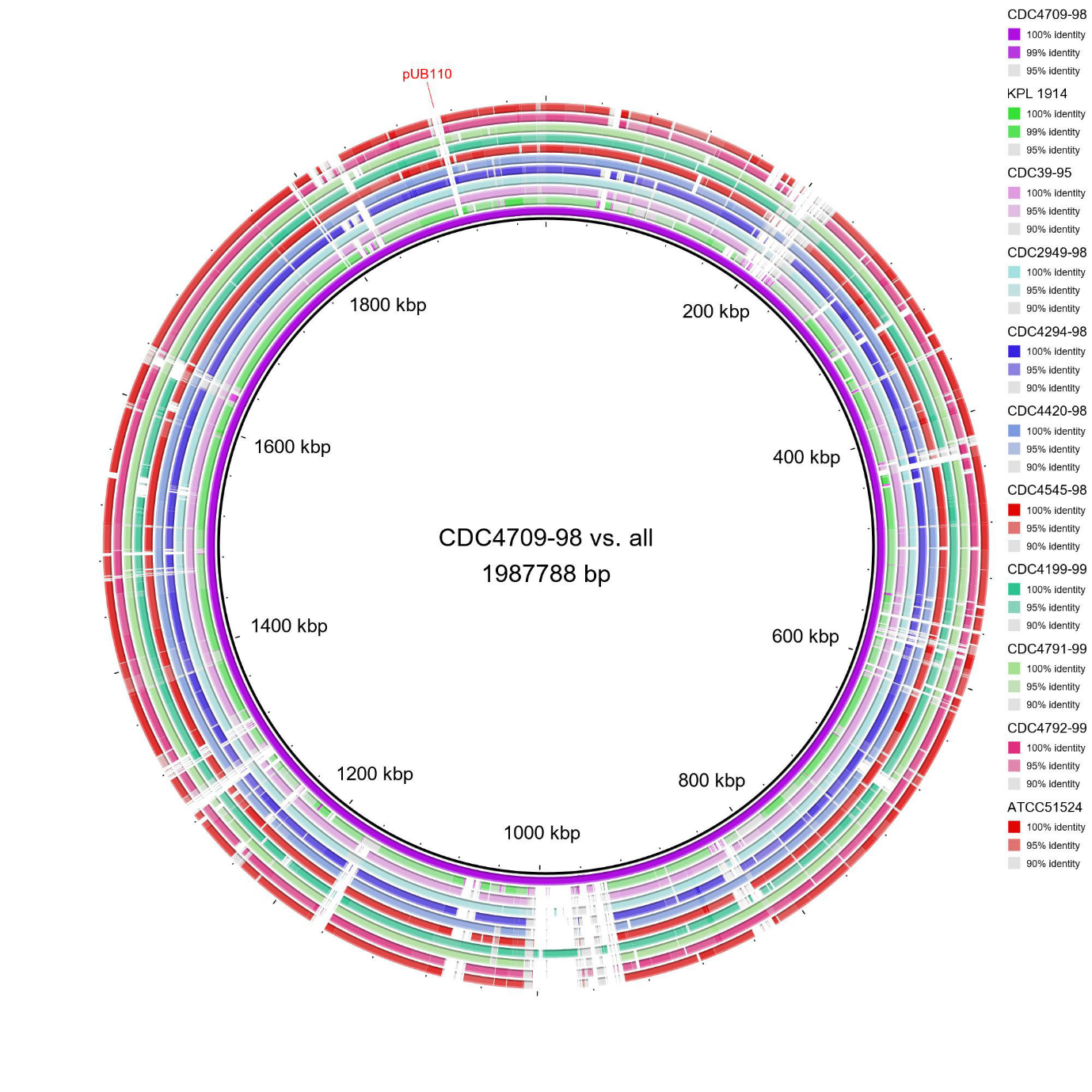

**Figure S7. Phylogenetic relationship between the 11 *D. pigrum* strains.** D. pigrum phylogenetic trees were constructed by using concatenated alignments of the intersection of predicted core genes (BDBH, COG, and OMCL, see above). Amino acids were aligned using Clustal Omega V.1.2.4 [7] and a maximum likelihood phylogenetic tree was generated using the LG model of amino-acid replacement matrix [8] as selected by smart model selection with Akaike information criterion [9] and 100 bootstrap replicates for branch support with PhyML (phylogenetic maximal likelihood) V. 3.0 [10] and visualized using FigTree V.1.4.4. A BIONJ distance-based tree was used as a starting tree to be redefined by the maximum likelihood algorithm. Bootstrap support values from 100 replicates are indicated above the branches. (**A**) Core genome AA tree of the 11 *D. pigrum* genomes. (**B**) Core genome AA tree with Alloiococcus otitis ATCC 51267 as an outgroup. Core genome size decreased to 866 clusters when including *A. otitis*.

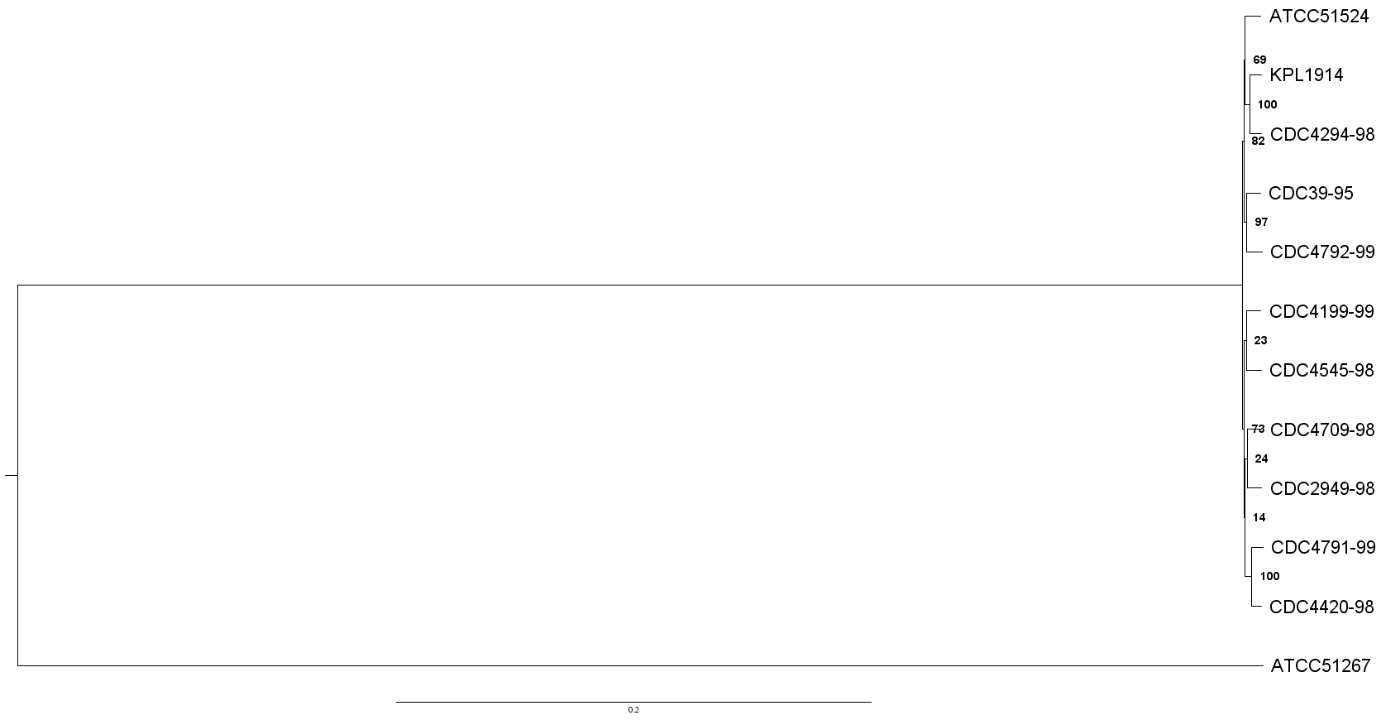
**7A**
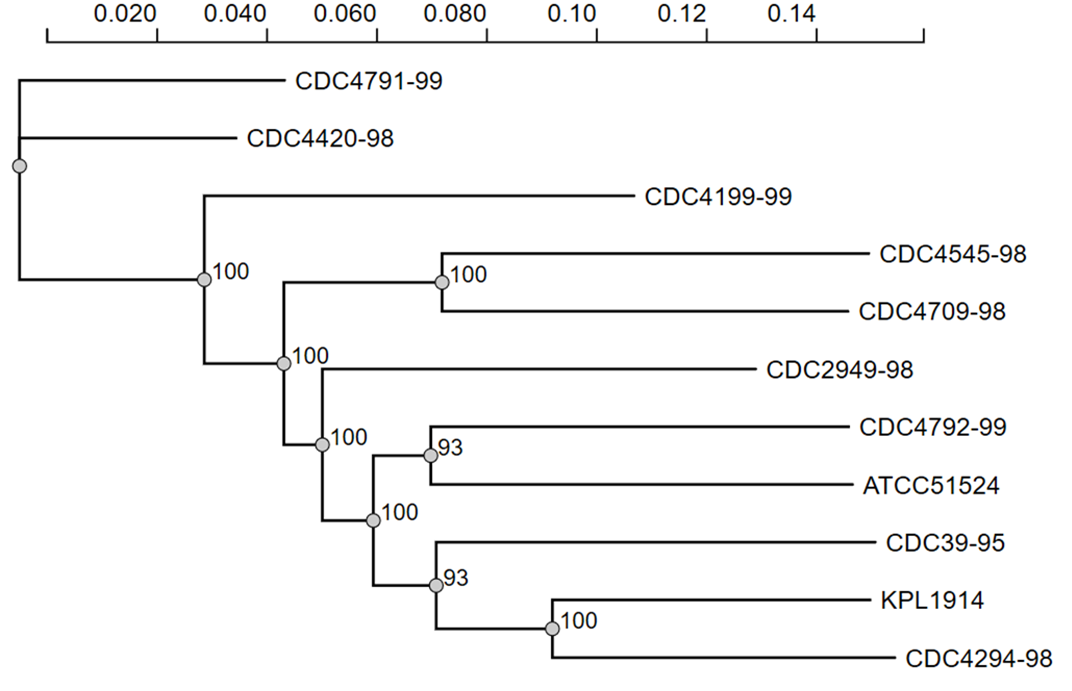
**7B**

**Table S5. Predicted biosynthetic gene clusters / secondary metabolites regions (antiSMASH version 5.0.0beta1-4e548fe)** [11]**.**

| **Strain** | **Bacteriocins** | **Lanthipeptides** |
| --- | --- | --- |
| KPL1914 | 2 | 1 |
| 39-95 | 1 | 1 |
| 2949-98 | 1 | 0 |
| 4294-98 | 2 | 0 |
| 4420-98 | 0 | 0 |
| 4545-98 | 1 | 0 |
| 4709-98 | 2 | 1 |
| 4199-99 | 3 | 0 |
| 4791-99 | 0 | 0 |
| 4792-99 | 2 | 0 |
| ATCC51524 | 0 | 2 |

**Figure S8. Predicted biosynthetic gene clusters / secondary metabolites regions (antiSMASH version 5.0.0beta1-4e548fe)** [11]**.**

**KPL1914**

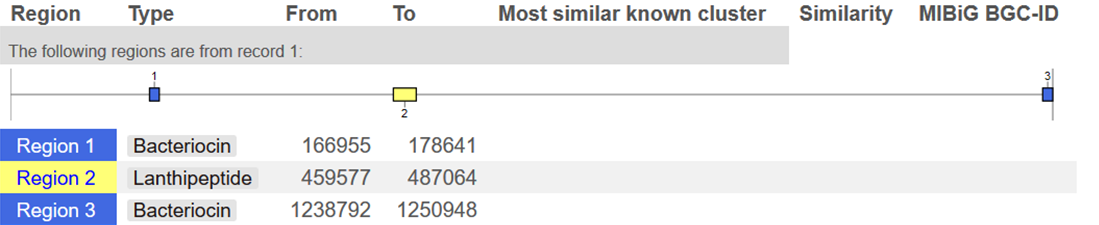

**CDC4709-98**

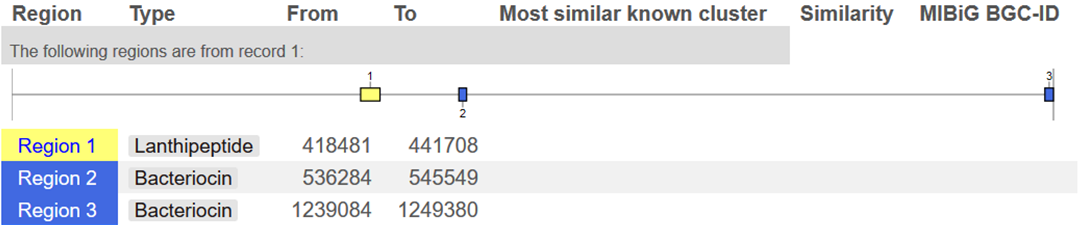

**CDC39-95**

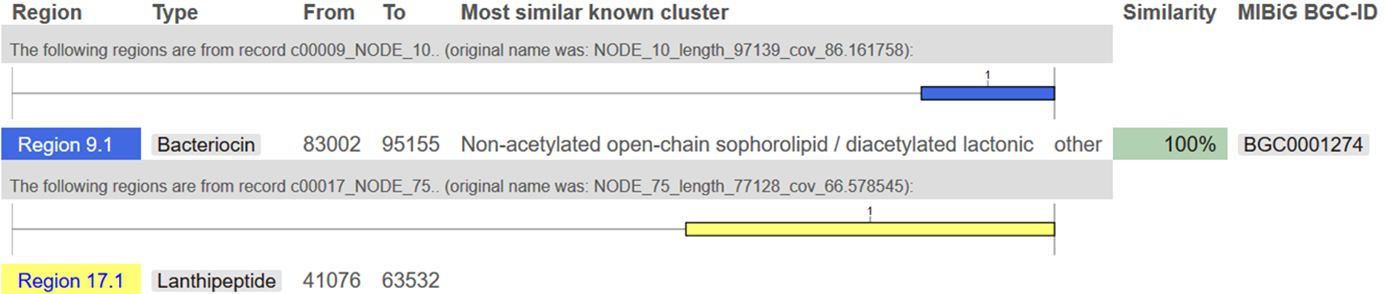

**CDC2949-98**

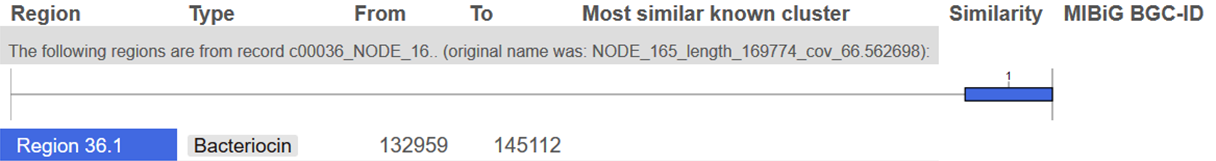

**CDC4294-98**

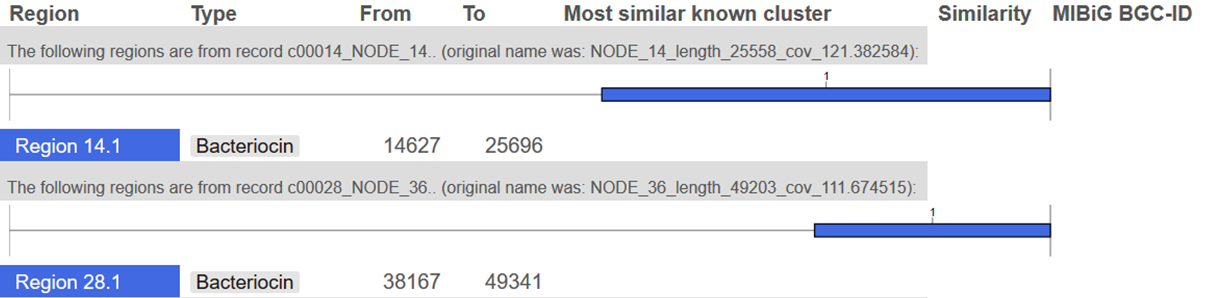

**CDC4545-98**

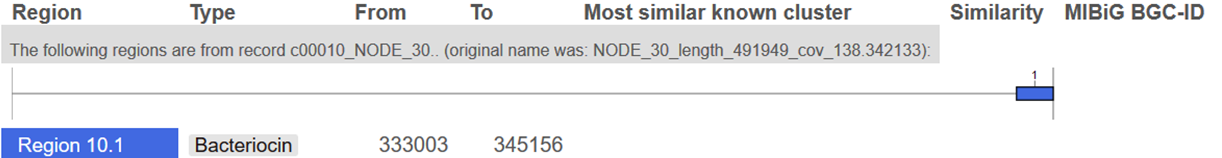

**CDC4199-99**

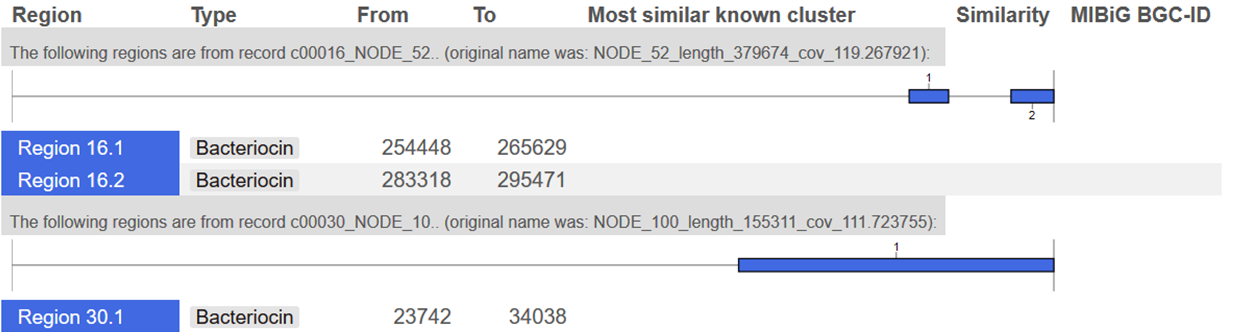

**CDC4792-99**

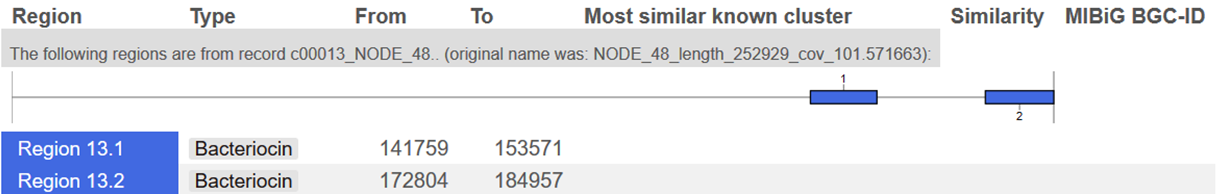

**ATCC51524**

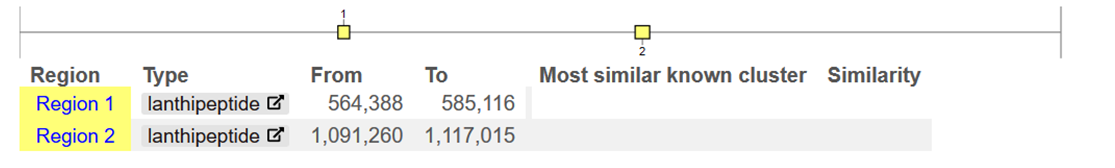

**Table S6. Species-level reanalysis of a pediatric nostril microbiota dataset.** (see excel file)

**Table S7: Non-*D. pigrum* bacterial strains used in this study**

| **Species** | **Strain** | **Internal Reference** | **Reference** | **Characteristics** |
| --- | --- | --- | --- | --- |
| *C. accolens* | KPL1818 | KPL1818 | [12] | primary adult human nostril isolate; lipid dependent |
| *C. pseudodiphtheriticum* | KPL1989 | KPL1989 | This study | primary adult human nostril isolate;  lipid independent |
| *C. pseudodiphtheriticum* | DSM44287^T^ | KPL2589 | [13] | type strain;  lipid independent |
| *C. propinquum* | DSM44285^T^ | KPL1955 | [14] | type strain;  lipid independent |
| *S. aureus* | Newman | KPL2023 | [15] | lab-adapted strain |
| *S. aureus* | JE2 | KPL2115 | [16] | plasmid-free derivative of USA300 LAC |
| *S. pneumoniae* | TIGR4 | KPL1904 | [17] | Clinical isolate |
| *S. pneumoniae* | DBL5 | KPL1905 | [18] | Clinical isolate |
| *S. pneumoniae* | 603 | KPL1906 | [19] | Clinical isolate |
| *S. pneumoniae* | WU2 | KPL1907 | [20] | Clinical isolate |

**References**

1.	Laclaire L, Facklam R: **Antimicrobial susceptibility and clinical sources of Dolosigranulum pigrum cultures.** *Antimicrob Agents Chemother* 2000, **44:**2001-2003.

2.	Aguirre M, Morrison D, Cookson BD, Gay FW, Collins MD: **Phenotypic and phylogenetic characterization of some Gemella-like organisms from human infections: description of Dolosigranulum pigrum gen. nov., sp. nov.** *J Appl Bacteriol* 1993, **75:**608-612.

3.	Contreras-Moreira B, Vinuesa P: **GET_HOMOLOGUES, a versatile software package for scalable and robust microbial pangenome analysis.** *Appl Environ Microbiol* 2013, **79:**7696-7701.

4.	Kaas RS, Friis C, Ussery DW, Aarestrup FM: **Estimating variation within the genes and inferring the phylogeny of 186 sequenced diverse Escherichia coli genomes.** *BMC Genomics* 2012, **13:**577.

5.	Darling AE, Mau B, Perna NT: **progressiveMauve: multiple genome alignment with gene gain, loss and rearrangement.** *PLoS One* 2010, **5:**e11147.

6.	Alikhan NF, Petty NK, Ben Zakour NL, Beatson SA: **BLAST Ring Image Generator (BRIG): simple prokaryote genome comparisons.** *BMC Genomics* 2011, **12:**402.

7.	Sievers F, Wilm A, Dineen D, Gibson TJ, Karplus K, Li W, Lopez R, McWilliam H, Remmert M, Soding J, et al: **Fast, scalable generation of high-quality protein multiple sequence alignments using Clustal Omega.** *Mol Syst Biol* 2011, **7:**539.

8.	Le SQ, Gascuel O: **An improved general amino acid replacement matrix.** *Mol Biol Evol* 2008, **25:**1307-1320.

9.	Lefort V, Longueville JE, Gascuel O: **SMS: Smart Model Selection in PhyML.** *Mol Biol Evol* 2017, **34:**2422-2424.

10.	Guindon S, Dufayard JF, Lefort V, Anisimova M, Hordijk W, Gascuel O: **New algorithms and methods to estimate maximum-likelihood phylogenies: assessing the performance of PhyML 3.0.** *Syst Biol* 2010, **59:**307-321.

11.	Blin K, Shaw S, Steinke K, Villebro R, Ziemert N, Lee SY, Medema MH, Weber T: **antiSMASH 5.0: updates to the secondary metabolite genome mining pipeline.** *Nucleic Acids Res* 2019.

12.	Bomar L, Brugger SD, Yost BH, Davies SS, Lemon KP: **Corynebacterium accolens Releases Antipneumococcal Free Fatty Acids from Human Nostril and Skin Surface Triacylglycerols.** *MBio* 2016, **7:**e01725-01715.

13.	Lehmann KB, Neumann RO: **Corynebacterium pseudodiphtheriticum.** In *Atlas und Grundriss der Bakteriologie und Lehrbuch der Speziellen Bakteriologischen Diagnostik.* 1920 edition. Munich: J.F. Lehmann; 1896: 571-572

14.	Riegel P, de Briel D, Prevost G, Jehl F, Monteil H: **Proposal of *Corynebacterium propinquum* sp. nov. for *Corynebacterium* group ANF-3 strains.** *FEMS Microbiology Letters* 1993, **113:**229-234.

15.	Miller KD, Hetrick DL, Bielefeldt DJ: **Production and properties of *Staphylococcus aureus* (strain Newman D2C) with uniform clumping factor activity.** *Thromb Res* 1977, **10:**203-211.

16.	Fey PD, Endres JL, Yajjala VK, Widhelm TJ, Boissy RJ, Bose JL, Bayles KW: **A genetic resource for rapid and comprehensive phenotype screening of nonessential *Staphylococcus aureus* genes.** *MBio* 2013, **4:**e00537-00512.

17.	Tettelin H, Nelson KE, Paulsen IT, Eisen JA, Read TD, Peterson S, Heidelberg J, DeBoy RT, Haft DH, Dodson RJ, et al: **Complete genome sequence of a virulent isolate of Streptococcus pneumoniae.** *Science* 2001, **293:**498-506.

18.	Lu YJ, Leite L, Goncalves VM, Dias Wde O, Liberman C, Fratelli F, Alderson M, Tate A, Maisonneuve JF, Robertson G, et al: **GMP-grade pneumococcal whole-cell vaccine injected subcutaneously protects mice from nasopharyngeal colonization and fatal aspiration-sepsis.** *Vaccine* 2010, **28:**7468-7475.

19.	Malley R, Lipsitch M, Stack A, Saladino R, Fleisher G, Pelton S, Thompson C, Briles D, Anderson P: **Intranasal immunization with killed unencapsulated whole cells prevents colonization and invasive disease by capsulated pneumococci.** *Infect Immun* 2001, **69:**4870-4873.

20.	Briles DE, Nahm M, Schroer K, Davie J, Baker P, Kearney J, Barletta R: **Antiphosphocholine antibodies found in normal mouse serum are protective against intravenous infection with type 3 streptococcus pneumoniae.** *J Exp Med* 1981, **153:**694-705.
