## SupplementalText for "*Dolosigranulum pigrum* cooperation and competition in human nasal microbiota"

**Supplemental Text**

### I. The core genome, chromosomal structure and phylogeny of *D. pigrum*

**The core genome of *D. pigrum* shows a high degree of sequence conservation**. From the nucleotide sequences of the 11 *D. pigrum* strains, an estimate of the upper bound of a conservative core genome for *D. pigrum* is 1200 coding sequencing (CDS) based on the overlap of three ortholog prediction algorithms (**Figure S1**). Two of these algorithms together estimated a lower bound of 1513 CDS for the *D. pigrum* accessory genome within a pangenome of 2729 CDS (based on an estimated core of 1216 CDS) (**Figure S2** and **Table S5**). Similarity matrices of the core genes and proteins from the 11 *D. pigrum* strains revealed a high degree of nucleotide (≥ 97%) and amino acid (≥ 96.8%) sequence conservation (**Figure S3A** and **S3B**, respectively).

To gain insight into the chromosomal structure of *D. pigrum*, we closed the genomes of two strains, CDC 4709-98 and KPL1914 using SMRT sequencing. MAUVE alignment revealed large blocks of synteny between these two closed genomes (**Figure S4**). Synteny analysis of the bidirectional best-hits core proteins from the RAST annotation revealed 1143 syntenic clusters (of 1206 total BDBH clusters). Ring plots using the closed genome of CDC 4709-98 as the reference (**Figure S5A**), because it is 5% (97 kb) larger than that of KPL1914 (**Figure S5B**), showed discrete areas of sequence absence in KPL1914 (**Figure S5C**) along with large areas of conservation. Similar areas of absence were visible when the other 9 strains were also compared to CDC 4709-98 (**Figure S5D**).

**A core-genome phylogeny reveals relationships between the 11 *D. pigrum* strains.** In 16S rRNA gene-based phylogenies, *D. pigrum* clades with other lactic acid bacteria and shares the closest node with *Alloiococcus otitis* (previously designated *A. otitidis*); both species are within the family *Carnobacteriaceae* (order *Lactobacillales*) [1]. Genomic content also identified *A. otitis* as its closest genome-sequenced neighbor by RAST. Using *A. otitis* ATCC 51267 as an outgroup, a core-genome, maximum likelihood-based phylogeny revealed a monophyletic *D. pigrum* clade (**Figure S6**). Although the 11 strains were isolated from different individuals in different geographic regions, mostly in the late 1990s (**Table S3**), there was a relatively small evolutionary distance. We next analyzed the predicted functions encoded in these *D. pigrum* genomes.

### II. Predicted biosynthesis, uptake and degradation by *D. pigrum* of amino acids, carbohydrates, polyamines and enzyme co-factors

**Methionine auxotrophy and degradation.** All 11 *D. pigrum* genomes lacked a complete known pathway for methionine biosynthesis. Methionine has been reported to be a limiting nutrient in the nasal passages of humans and its synthesis is upregulated in *S. aureus* growing in synthetic nasal medium [2]. In contrast, all 11 encoded two full sets of the genes required for methionine degradation in different regions on their chromosome suggesting external dependence. Each set includes *metN* (methionine ABC transporter ATP-binding protein), *metP* (methionine ABC transporter permease protein), *metQ* (methionine ABC transporter substrate-binding protein) and *metT* (methionine transporter).

**Arginine auxotrophy and degradation.** All 11 *D. pigrum* strains lacked most of the genes required for synthesis of arginine from glutamate (e.g., *argB, argC, argD, argF, argG* and *argH*) suggesting likely auxotrophy. All 11 strains contained the genes of the arginine deiminase pathway (*arcA, arcB, arcC, arcD* and *arcR*) suggesting *D. pigrum* may uptake extracellular arginine.

**Glutamine.** Glutamine synthetase I (EC 6.3.1.2) was predicted in all 11 strains and catalyzes the reaction of L-glutamine from L-glutamate and NH_4_^+^ using ATP. Glutamate racemase (EC 5.1.1.3) was predicted in all 11 and catalyzes D- / L-glutamate interconversion. NAD(P)-specific glutamate dehydrogenase (EC 1.1.1.4) GdhA was predicted in all 11 and catalyzes the formation of L-glutamate and H_2_O from 2-oxoglutarate, ammonia and NADH+H^+^ and vice versa.

**Polyamine auxotrophy and transport.** The absence of predicted genes required for *de novo* synthesis of the essential polyamines putrescine and spermidine indicates that *D. pigrum* likely relies on extracellular polyamines. Consistent with this, all strains harbor putative ABC type spermidine transporter components (SPERta, SPERtb, SPERtc, SPERtd) adjacent to each other as well as putative putrescine transporter genes (*potA*, *potB*, *potC*, and *potD*).

**Biotin auxotrophy and transport.** A biotin uptake system was predicted in all 11 *D. pigrum* strains (Substrate-specific component BioY of biotin ECF transporter) and no biosynthesis was predicted. Biotin-protein ligase (EC 6.3.4.15) was predicted only in KPL1914 and ATCC51524.

**Nicotinic acid (niacin) auxotrophy and transport.** NAD and NADP are critical factors for cellular metabolism**.** However, *D. pigrum* lacked known genes for *de novo* synthesis and for salvage of nicotinate or nicotinamide. However, the presence of *niaP* predicts uptake of nicotinic acid (niacin) by all 11 strains. All strains also encoded genes needed to convert niacin or nicotinamide to NAD+ and NADP: nicotinamidase (EC 3.5.1.19), nicotinate phosphoribosyltransferase (EC 2.4.2.11), nicotinate-nucleotide adenylyltransferase (EC 2.7.7.18), NAD synthetase (EC 6.3.1.5), NAD kinase (EC 2.7.1.23) and NadR transcriptional regulator.

***D. pigrum* encodes mechanisms for acquiring essential metal cofactors from the host environment.** The nasal environment is low on essential metal ions such as iron, zinc and manganese and host metal sequestration using lactoferrin and calprotectin is an important defense mechanism against bacterial growth. Bacteria have acquired mechanisms to escape this nutritional immunity [2, 3]. We therefore searched for genes predicted to encode for siderophores and transporters for heme, manganese and zinc. All 11 *D. pigrum* genomes harbored a predicted iron compound ABC uptake transporter ATP-binding protein (hemin uptake system subsystem) and a manganese ABC-type transporter. Additionally, six of the CDC strains (4294-98, 4420-98, 4545-98, 4199-99, 4791-99, 4792-99) had predicted ferric iron ABC transporter and/or iron compound ABC uptake transporter genes.

### III. Predicted carbohydrate metabolism by *D. pigrum* via homofermentation to lactate

**There is no tricarboxylic acid (TCA) cycle present in *D. pigrum***. Only fumarate-hydratase (EC 4.2.1.2) and TCA associated dihydrolipoyl dehydrogenase (EC 1.8.1.4) were predicted in all *D. pigrum* genomes.

**Anaerobic respiratory reductases.** We did not identify butyryl-CoA-reductase (EC 1.3.8.1) or any other predicted anaerobic reductases in all *D. pigrum* strains. However, *D. pigrum* CDC 4545-98 encoded an arsenate reductase (EC 1.20.4.1).

**Identification of a V-type ATPase in all 11 isolates**. V-ATPases hydrolyse ATP to drive a proton pump but cannot work in reverse to synthesize ATP.

**Glycolysis (Embden-Meyerhof-Parnas pathway, EMP).** Enzymes present in all 11 isolates included glucokinase, glucose-6-phosphate isomerase, 6-phosphofructokinase, fructose-bisphosphate aldolase class II-1,6-bisphosphate-aldolase, triose phosphate isomerase, NAD-dependent glyceraldehyde-3-phosphate dehydrogenase, phosphoglycerate kinase, 2,3-bisphosphoglycerate-independent phosphoglycerate mutase, enolase and pyruvate kinase. Three strains (KPL1914, CDC 39-95, CDC 4792-99) also encoded a predicted fructose-bisphosphate aldolase class I (EC 4.1.2.13), whereas all strains harbored a predicted triosephosphate isomerase.

As noted in the main text, all 11 strains encoded a predicted L-lactate-dehydrogenase (EC 1.1.1.27), which catalyzes the reduction of pyruvate to lactate regenerating NAD+ for glycolysis (GAPDH step), consistent with homofermentation to L-lactate as the primary product of glycolysis. Moreover, the absence of xylulose-5-phosphate phosphoketolase (EC 4.1.2.9) is inconsistent with (obligate) heterofermentation, since bacteria that heteroferment lack aldolase and have to shunt through the pentose phosphate or phosphoketolase pathway [4]. In addition to homofermentation to lactate, some end product flexibility, which is probably produced under differing environmental/nutritional conditions, is predicted by the presence in all 11 genomes of genes needed to accomplish mixed-acid fermentation with potential production of formate, acetate and ethanol (i.e., enzymes pyruvate formate-lyase (EC 2.3.1.54), phosphate acetyltransferase (EC 2.3.1.8) and acetate kinase (EC 2.7.2.1), as well as acetaldehyde dehydrogenase (1.2.1.10) and alcohol dehydrogenase (EC 1.1.1.1)).

**Sialidases.** The original species description of *D. pigrum* reports production of acid from D-glucose, galactose, D-fructose, D-mannose, maltose and L-fucose in two isolates with strain-level variation in producing acid from D-arabinose, mannitol, sucrose and N-acetyl-glucosamine [5]. Glucose is the main monosaccharide detected in the nasal environment [2]. Host-cell-surface- and host-mucin-derived sialic acids are another important potential source of monosaccharides and all 11 *D. pigrum* genomes harbor a putative sialidase as well as a predicted transporter (sodium solute symporter) and catabolic enzymes suggesting it utilizes sialic acid in the nasal passages.

### IV. *D. pigrum* is predicted to be broadly susceptible to antibiotics

**Antibiotic resistance prediction.** We analyzed all 11 *D. pigrum* genomes for putative antibiotic resistance genes and mutations that confer antibiotic resistance using the Resistance Gene Identifier (RGI) on the Comprehensive Antibiotic Resistance Database (CARD) [6] allowing only perfect and strict results. This identified a candidate in only one strain. These data are consistent with the report from LaClaire and Facklam of *D. pigrum* susceptibility to amoxicillin, cefotaxime, cefuroxime, clindamycin, levofloxacin, meropenem, penicillin, quinupristin-dalfopristin, rifampin, tetracycline, and vancomycin for all tested *D. pigrum* strains [7]. *D. pigrum* strain CDC 4709-98 alone encoded a predicted bleomycin resistance protein (BRP), which was predicted with 100% amino acid in a protein homolog model. In agreement, ResFinder [8] identified kanamycin nucleotidyltransferase (*aadD*, aka ANT(4')-Ia, aminoglycoside adenyltransferase AAD, spectinomycin resistance; streptomycin resistance; transferase) in this strain with 99.74 % sequence identity. Detailed analysis of this region with PlasmidFinder [9] and BLAST [10] revealed loci with sequence identity to portions of plasmid pUB110 from *S. aureus* [11] suggesting integration of, or at least part of, this plasmid into the genome of strain CDC 4709-98.

**References**

1.	Yarza P, Richter M, Peplies J, Euzeby J, Amann R, Schleifer KH, Ludwig W, Glockner FO, Rossello-Mora R: **The All-Species Living Tree project: a 16S rRNA-based phylogenetic tree of all sequenced type strains.** *Syst Appl Microbiol* 2008, **31:**241-250.

2.	Krismer B, Liebeke M, Janek D, Nega M, Rautenberg M, Hornig G, Unger C, Weidenmaier C, Lalk M, Peschel A: **Nutrient limitation governs Staphylococcus aureus metabolism and niche adaptation in the human nose.** *PLoS Pathog* 2014, **10:**e1003862.

3.	Krismer B, Weidenmaier C, Zipperer A, Peschel A: **The commensal lifestyle of Staphylococcus aureus and its interactions with the nasal microbiota.** *Nat Rev Microbiol* 2017, **15:**675-687.

4.	Buyze G, Van Den Hamer CJ, De Haan PG: **Correlation between hexosemonophosphate shunt, glycolytic system and fermentation-type in Lactobacilli.** *Antonie Van Leeuwenhoek* 1957, **23:**345-350.

5.	Aguirre M, Morrison D, Cookson BD, Gay FW, Collins MD: **Phenotypic and phylogenetic characterization of some Gemella-like organisms from human infections: description of Dolosigranulum pigrum gen. nov., sp. nov.** *J Appl Bacteriol* 1993, **75:**608-612.

6.	McArthur AG, Waglechner N, Nizam F, Yan A, Azad MA, Baylay AJ, Bhullar K, Canova MJ, De Pascale G, Ejim L, et al: **The comprehensive antibiotic resistance database.** *Antimicrob Agents Chemother* 2013, **57:**3348-3357.

7.	Laclaire L, Facklam R: **Antimicrobial susceptibility and clinical sources of Dolosigranulum pigrum cultures.** *Antimicrob Agents Chemother* 2000, **44:**2001-2003.

8.	Zankari E, Hasman H, Cosentino S, Vestergaard M, Rasmussen S, Lund O, Aarestrup FM, Larsen MV: **Identification of acquired antimicrobial resistance genes.** *J Antimicrob Chemother* 2012, **67:**2640-2644.

9.	Carattoli A, Zankari E, Garcia-Fernandez A, Voldby Larsen M, Lund O, Villa L, Moller Aarestrup F, Hasman H: **In silico detection and typing of plasmids using PlasmidFinder and plasmid multilocus sequence typing.** *Antimicrob Agents Chemother* 2014, **58:**3895-3903.

10.	Altschul SF, Gish W, Miller W, Myers EW, Lipman DJ: **Basic local alignment search tool.** *J Mol Biol* 1990, **215:**403-410.

11.	McKenzie T, Hoshino T, Tanaka T, Sueoka N: **The nucleotide sequence of pUB110: some salient features in relation to replication and its regulation.** *Plasmid* 1986, **15:**93-103.
